# Cardiolipin-mimic lipid nanoparticles without antibody modification delivered senolytic in-vivo CAR-T therapy for inflamm-aging

**DOI:** 10.1101/2024.11.21.624667

**Authors:** Zihan Zhang, Bin Ma, Buyao Li, Zhiwei Li, Min Gao, Hailong Zhao, Rui Peng, Jiang Hu, Yu Wang, Wei You, Xun Gui, Rui Wang, Xiaoqing Hu, Beidi Chen, Yuanjie Zhang, Yanyun Hao, Demin Zhou, Yun Yang, Mi Deng, Lei Miao

## Abstract

mRNA-based in vivo CAR T cell engineering offers advantages over ex vivo therapies, including streamlined manufacturing and transient expression. However, current delivery requires antibody-modified vehicles with manufacturing challenges. In this study, inspired by cardiolipin, we identified a cardiolipin-like di-phosphoramide lipid that improved T cell transfection without targeting ligands, both in vivo and in vitro. The T cell-favored tropism is likely due to the lipid’s packing, shape, and rigidity. Encapsulating circular RNA further prolonged mRNA expression in the spleen and T cells. Using PL40 lipid nanoparticles, we delivered mRNA encoding a CAR targeting the senolytic and inflammatory antigen urokinase-type plasminogen activator receptor (uPAR), alleviating uPAR-related liver fibrosis and rheumatoid arthritis (RA). Single cell sequencing in humans confirmed uPAR’s relevance to senescence and inflammation in RA. To further enhance clinical translation, we screened and humanized scFvs against uPAR, establishing PL40 mRNA encoding a circular human uPAR CAR, with potential for treating aging-inflamed disorders.

**One Sentence Summary:** We’ve developed a unique class of Cardiolipin-mimic lipids that facilitate mRNA delivery to T cells in vivo without the need for antibody modification, enhancing the treatment of liver fibrosis and rheumatoid arthritis through circular CAR uPAR RNA and propelling the clinical application of humanized CAR against human uPAR.

## INTRODUCTION

Over the past several decades, chimeric antigen receptor (CAR) T cell therapy has revolutionized the treatment of previously incurable hematological malignancies, including lymphoma, leukemia, and multiple myeloma (*1*). The remarkable specificity and potent cytotoxicity of CAR T cells have sparked interest in exploring non-cancer applications, though their clinical utility remains limited. A few case reports have described the employment of CD19-targeted CAR T cells to treat autoimmune disorders (*2, 3*). More recently, CAR T cells have shown promise in targeting cellular senescence, a hallmark of chronic inflammatory diseases (termed ‘inflamm-aging’) (*4*). Potential senolytic targets include the urokinase-type plasminogen activator receptor (uPAR) and Natural killer group 2 member D (NKG2D) ligands (*5–7*). While senolytic drugs have exhibited preclinical potential in various chronic diseases, the specific identification of senescent cell markers remains challenging. CAR T and macrophages offer advantages in this context, given their high specificity.

Compared to conventional *ex vivo* CAR T cell therapies, mRNA-based *in vivo* CAR T cell engineering offers several potential advantages for treating inflamm-aging. The *in vivo* approach eliminates the need for complex and costly *ex vivo* cell processing. Moreover, the transient nature of mRNA-based *in vivo* CAR T cells avoids the permanent genomic alterations seen with viral vector-based approaches (*8*). This may also help mitigate cytokine release syndrome and the on-target/off-tissue adverse events often observed with persistent activation of conventional CAR T cells (*9*). While the transient expression could be a drawback for solid cancer treatment, it may be a benefit for inflamm-aging where leukocyte access is not a hurdle. Overall, mRNA-based *in vivo* CAR T cell engineering presents compelling advantages over conventional approaches for treating inflamm-aging diseases.

Numerous studies have demonstrated the delivery of CAR mRNA to T cells using antibody-targeted lipid nanoparticles (tLNPs) or polymeric nanoparticles (*10–12*). While antibody targeting is well-established, the scale-up process poses challenges. There are concerns about batch-to-batch reproducibility, degree of antibody modification, and conjugation yield. Improper antibody selection may also exacerbate LNP clearance or lead to target cell over-activation (*12*). In contrast, non-targeted LNPs are easier to manufacture, potentially offering higher safety. Interestingly, some lipid formulations, like adamantanamine-derived cationic lipids (*13–15*), have demonstrated intrinsic tropism for T cells and B cells, suggesting the possibility of improved immune cell targeting without the need for antibody conjugation.

In the present study, we addressed *in vivo* T cell delivery challenges by drawing inspiration from the natural cardiolipin structure and designing novel cardiolipin-mimetic phosphoramide (CAMP) lipids. CAMP lipids were incorporated into LNPs to remodel the structure and enhance T cell delivery both *in vitro* and *in vivo*. Our findings indicate that the T cell-favored tropism of the CAMP-containing LNPs was associated with changes in their shape and rigidity.

Specifically, CAMP lipids increased LNP hardness and phase separation, improving cellular uptake by T cells. To further prolong the *in vivo* CAR protein expression, we modified the mRNA payload by converting it into a circular RNA format. This circular RNA approach was found to enhance splenic targeting and prolong mRNA expression. We utilized the CAMP-containing LNP system to encapsulate CAR against uPAR, that was expressed in monocytes/macrophages and senescent fibroblasts, demonstrating proof-of-concept for treating liver fibrosis and rheumatoid arthritis (RA). Additionally, we have laid the groundwork for the humanized design and screening of a CAR targeting the human uPAR antigen, which may facilitate the clinical translation of this *in vivo* CAR therapeutic approach for treating inflamm-aging symptoms.

## RESULTS

### Development of cardiolipin-mimic phosphoramide (CAMP) lipid library for RNA delivery to primary human T cells

Cardiolipin, a major component of the inner mitochondrial membrane, plays a crucial role in membrane folding and curvature formation (Fig. 1A) (*16, 17*). Considering that the lipid structure can influence the packing and behavior of lipid nanoparticles (LNPs) and their interaction with biological membranes such as endosomes, we decided to utilize cardiolipin as a starting point to develop a library of modified cationic lipids. We synthesized a library of 23 purified ionizable cationic lipids, termed Cardiolipin-Analog Phosphoramide (CAMP) lipids, by modifying the negatively charged biphosphate headgroup of cardiolipin (Fig. 1B). The CAMP library was synthesized using POCl3 as a starting material, with three sequential additions of nucleophilic reagents at controlled temperatures. The purified CAMP lipids were then formulated with dioleoylphosphatidylethanolamine (DOPE), cholesterol, and C14-PEG into LNP formulations. mRNA encoding firefly luciferase (mLuc) was encapsulated within the CAMP LNPs and applied to primary human T cells at a concentration of 60 ng/1×105 cells. After 24 hours, luciferase expression was assessed through luminescence measurement.

**Figure 1.**
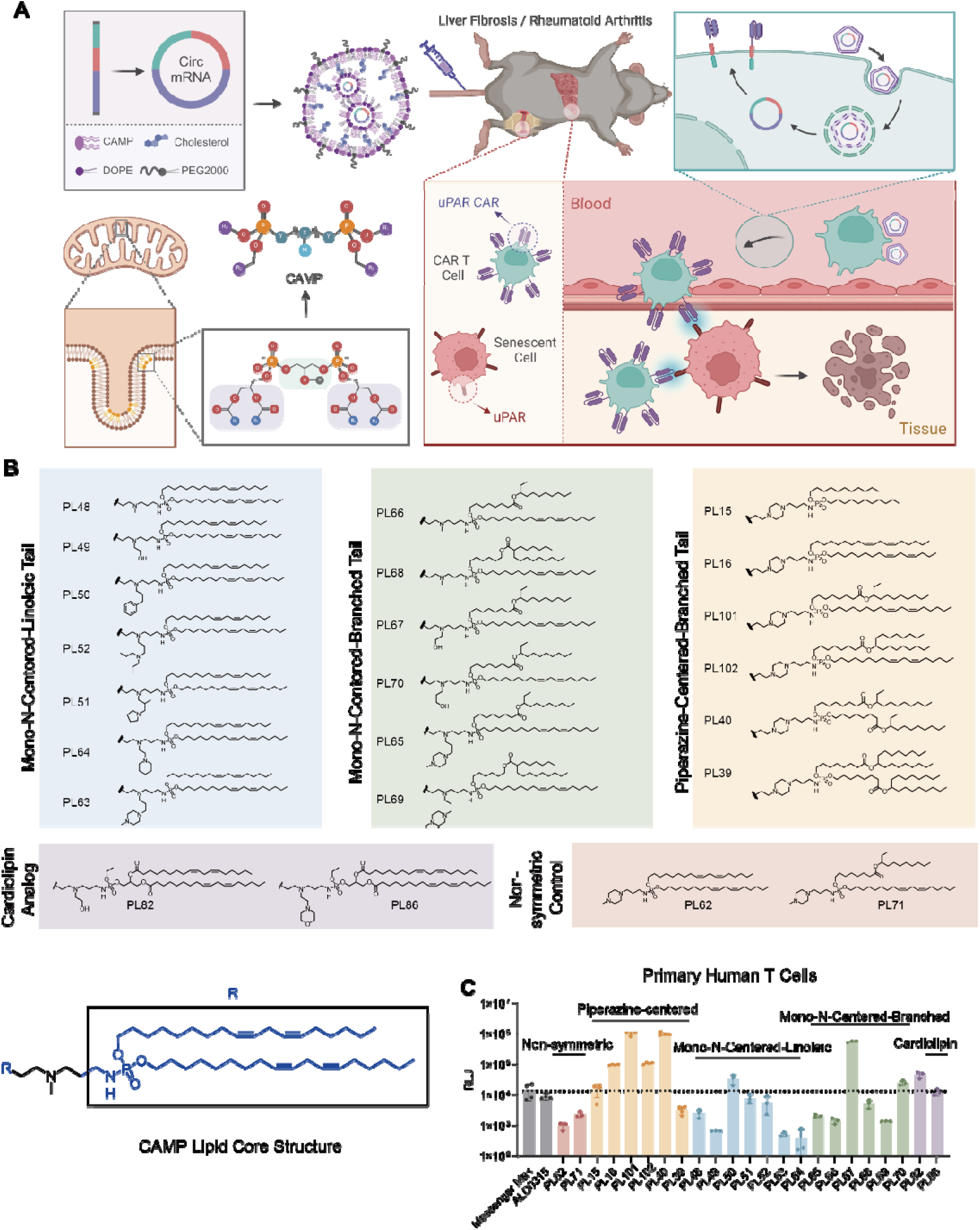
Development of CAMP lipid library for RNA delivery to primary T cells. **(A)** Schematic illustration of the similarity of T cell membrane structure and mitochondria inner membrane, with cardiolipin as the core lipid. Cardiolipin is used as the starting core structure to be modified into ionizable lipids. **(B)** The library of ionizable CAMP lipids. CAMP lipids are all lipids with symmetric structures. The core structure is presented. **(C)** Screening of the CAMP LNPs in primary human T cells. CAMP lipids were formulated with other helper lipids into LNPs. mLuc were delivered by CAMP LNPs at a dose of 60 ng mRNA/10^6^ cells. Luminescence was measured after 24 h (Messenger Max ∼ PL39, n = 4; PL48 ∼ PL86, n = 3; mean ± SD), “n” indicated biologically independent samples. The schematic in A was created with BioRender.com.

We found that the cardiolipin analogs PL82 and PL86, where the phospho-ester structure and glycerol linker were maintained but the central carbon was substituted with an amino group and the negative phosphate charge was acetylated, showed around a 10-fold increase in T cell transfection compared to the controls ALC0315 LNPs and Messenger Max (Fig. 1C). In contrast, replacing the glycerol linkage with two phospho-esters to simplify the symmetric structure did not improve T cell transfection. Interestingly, introducing a piperazine ring as the central core significantly enhanced T cell transfection (PL16/40/101/102) by 50-100 folds (Fig. 1C). Further improvements were observed by incorporating an ester bond with an asymmetric alpha-carbon ethyl branch (PL101 and PL40). This trend was also recapitulated in mono-N-centered CAMP lipids (PL67 and PL70). Ultimately, when compared to the MessengerMax and ALC0315 LNPs, the CAMP library identified seven LNPs that resulted in significantly higher luciferase expression in primary human T cells. Specifically, PL101, PL40, and PL67 LNPs exhibited over a 100-fold increase in expression compared to the positive controls (Fig. 1C).

### Structure and morphology impact PL40 CAMP LNP delivery of mRNAs to T cells

We then utilized the piperazine-centered CAMP lipids to transfect various types of cells (Fig. 2A). Since the transfection efficiencies were relatively low in NK92 cells, T cells remained one of the major cell types that preferentially expressed the mRNA cargo delivered by PL101 and PL40 LNPs (Fig. 2B, fig. S1A, D, E). These improvements were also observed in human T cells from additional donors and mouse primary T cells (Fig. 2C and fig. S1F).

**Figure 2.**
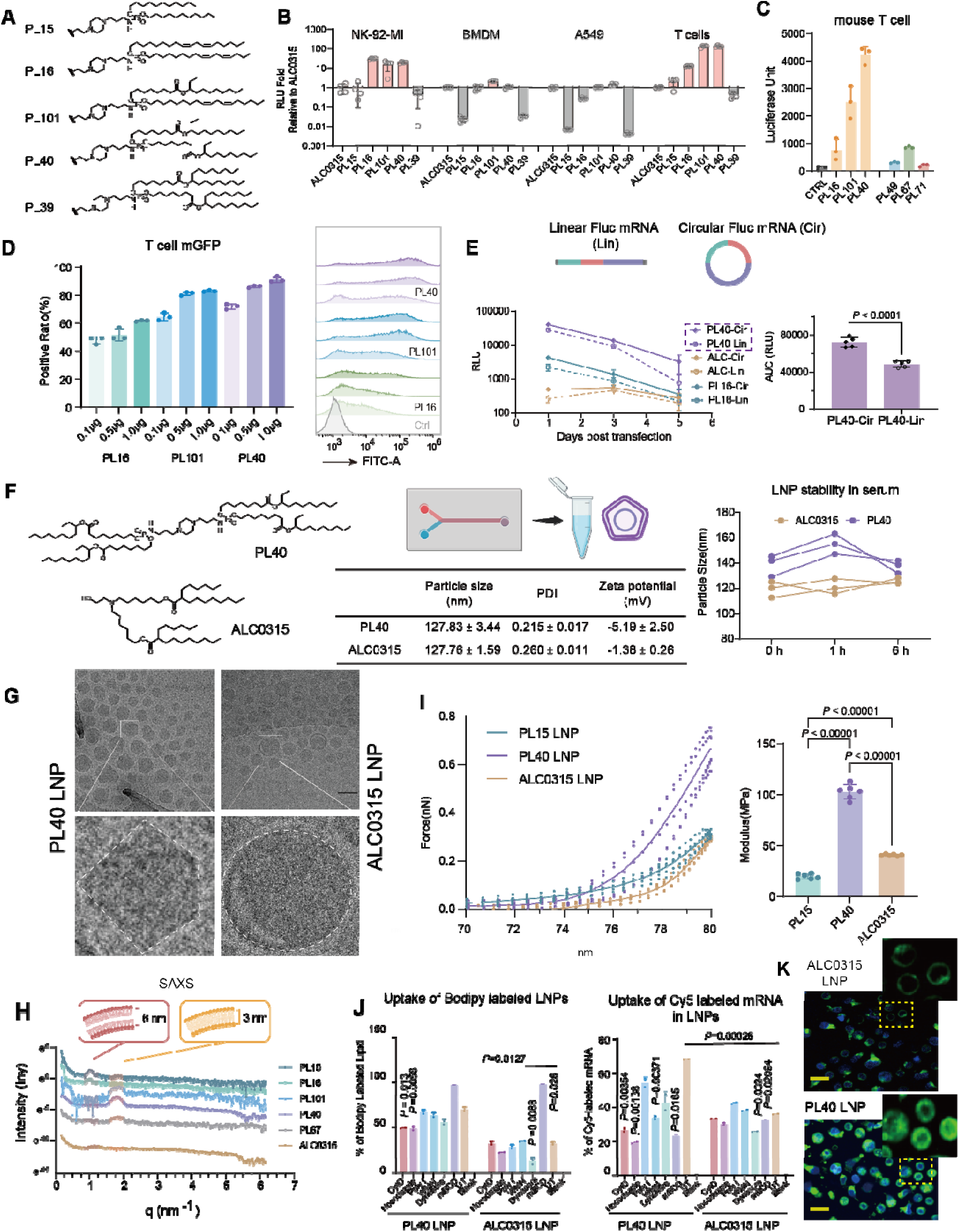
Structure and morphology impact PL40 CAMP LNP delivery of mRNAs to T cells. **(A)** The structure of CAMP lipids screened out for further study. **(B)** The transfection efficiency of CAMP lipid to various types of cells. The 96-well plate was paved with NK92mi, BMDM, A549 and human primary T cell respectively, 60 ng/well mLuc was transfected and measured at 24 h (BMDM n = 3; others n = 4; mean ± SD). **(C)** The transfection efficiency of CAMP lipids in primary mouse T cells. CAMP LNPs encapsulating mLuc were applied to transfect mouse primary T cell, at a dose of 60 ng/10^5^ cells. Luciferase intensity was measured at 24 h (n = 3, mean ± SD). **(D)** The % of T cells transfected with CAMP lipids encapsulating mGFP, 0.1 μg, 0.5 μg and 1 μg GFP mRNA were transfected to 10^5^ T cells (n = 3 per group per time point, mean ± SD). GFP^+^ cells were detected at 40 h by flow cytometry. **(E)** T cells were transfected with Circ and Lin mLuc encapsulated by ALC, PL40, PL16 LNPs. Luminescence was detected on day 1, 3, and 5, respectively (n = 5, mean ± SD). The AUC of PL40 was calculated. **(F)** Particle size, zeta potential and stability of PL40 and ALC0315 LNPs were detected (n = 3). **(G)** Representative Cryo-TEM images of PL40 and ALC0315 LNPs. **(H)** SAXS data of CAMP lipid LNPs and ALC0315 LNPs. Lamellar distance was calculated based on q value. **(I)** Representative AFM images, the force curve and Yang’s modulus of PL15, PL40, ALC0315 LNPs. Force curve and Yang’s modulus were measured by AFM (n = 5, mean ± SD). **(J)** Cellular uptake of ALC0315 and PL40 LNPs. T cells were transfected by Cy5 labeled mRNA encapsulated in Bodipy labeled LNPs at a dose of 0.5 μg/10^5^ cells. Small molecule inhibitors for multiple difference endocytosis pathways were added. % of LNP positive cells were detected by flow cytometry 4 h after transfection (n = **2,** mean ± SD). Experiments were repeated twice with similar results. (**K**) The bodipy labeled lipid (green) and Hoechst 33342 (blue) florescence images for illustration of LNP cellular uptake. The images were acquired by High Content Imaging and Analysis System (Cell Voyager CV8000, Yokogawa), images were taken from 3 independent samples, one representative was shown, scale bar indicated 10 µm. “n” indicated biologically independent samples. Statistical significance was calculated through two-tailed unpaired Student’s *t*-test (e), One-way ANOVA with Tukey test (i), Dunnett test (j).

To quantify the percentage of T cells expressing the mRNA, we encapsulated mRNA encoding GFP. All LNPs demonstrated dose-dependent GFP expression, with PL40 exhibiting the highest efficiency, allowing over 90% of cells to express the cargo at doses beyond 0.5 μg (Fig. 2D). Recognizing the enhanced stability and extended half-life of circular RNA (CircRNA) in HEK293T cells (fig. S2), we further utilized the CAMP library to encapsulate Circ mLuc (Fig. 2E). To generate CircRNA cargoes, we employed a previously reported Group II intron-based self-splicing method to produce the Circ mLuc, which was subsequently purified by FPLC to get pure Circ mLuc (fig. S3) (*18, 19*). In human primary T cells, Circ mLuc still prolonged the protein expression within five days as compared to the linear counterpart, highlighting the advantage of CircRNA for protein expression (Fig. 2E). Additionally, all lead CAMP formulations exhibited no observed toxicity to T cells at increased dosages (fig. S1B, C). Ultimately, PL40 was selected as the lead candidate. PL40 LNPs were characterized by >80% encapsulation efficiency, a particle size of approximately 120 nm, and zeta potential of –5.19 mV (Fig. 2F). Additionally, PL40 LNPs exhibited optimal plasma stability, similar to the commercial ALC0315 LNPs (Fig. 2F).

To investigate the mechanism underlying the improved transfection of PL40 in T cells, we examined the morphological and structural differences at the nanoscale between PL40 LNPs and the conventional ALC0315 LNPs using cryo-transmission electron microscopy (Cryo-TEM) (Fig. 2G). ALC0315 exhibited a uniform lamellar curvature on the surface and a relatively amorphous particle core. In contrast, PL40 displayed a highly faceted surface with a thinner lamellar structure and a denser particle core. This multiple faceted surface morphology has been observed previously in liposomes and LNPs containing beta-sitosterol (*20*). It is likely the result of phase separation of different lipid domains. Phase separation is also a mechanism by which cardiolipin forms the cristae of the mitochondrial inner membrane (*21, 22*). At the phase boundaries, a higher degree of disorder and packing defects are prevalent due to the mismatch in spontaneous curvature between different domains(*23–25*). These defects could potentially facilitate the fusion of LNPs with membranes, particularly in T cells where the membrane may also exhibit phase-separated defects.

To further investigate the internal structure of LNPs, we conducted small-angle X-ray scattering (SAXS) experiments (Fig. 2H). The results showed that LNPs with lower T cell expression, such as ALC0315, PL15, and PL16, exhibited a peak at q∼1 nm^-1^, a characteristic of an organized lamellar phase with mRNA either associated with one bilayer or sandwiched between bilayers. The lamellar spacing, calculated as d = 2π/q_m_, was estimated to be 6.64 nm for all three LNPs. In contrast, PL40, PL101, and PL67, which showed enhanced mRNA expression in T cells, presented a peak at q∼2 nm^-1^, suggesting a smaller d-spacing (∼3 nm) (*26*). This smaller d-spacing may result from tighter packing of lipids and mRNA, which is consistent with the denser lamellar and particle structure observed in Cryo-TEM. Next, we investigated whether the change in morphology and structure would impact the mechanical rigidity of the lipid membrane.

Atomic force microscopy (AFM) was used to evaluate the bending modulus, one key parameter defining rigidity, of LNPs in an aqueous phase (Fig. 2I) (*27*). The slope of the curve for PL40 was much steeper than that of ALC0315 and PL15 LNPs, indicating a significantly larger bending modulus for PL40 LNPs. The unique surface morphology and higher membrane bending modulus of PL40 LNPs contributed to improved cellular uptake in T cells. By modifying the LNPs with either dye labelled lipid or mRNA, we showed that PL40 exhibited ∼2-fold increase in cellular uptake in primary T cells, as compared to ALC0315 LNPs, which were impacted by actin orientations and micropinocytosis (*28*) (using inhibitors such as Cytochalasin D, CytD and Nocodazole, Fig. 2J and K). Besides uptake, the multi-chain core structure is also likely to facilitate membrane fusion and endosomal release once protonated, as previously mentioned (*29*). Taken together, the morphology and structure of PL40 are one of the reasons that facilitate its endocytosis and endosomal escape in T cells (Fig. 2K).

### PL40 LNPs delivered mRNA to T cells in vivo

Next, we investigated the *in vivo* delivery of mRNA to T cells using CAMP LNPs. We intravenously (*i.v.*) administered ALC0315, PL101 and PL40 LNPs carrying either linear (Lin) or Circ mLuc to C57 mice. Mice were sacrificed at 6 and 24 h post injection, and the bioluminescence of their organs were examined (Fig. 3A, fig. S4). Consistent with previous reports, the ALC0315 LNPs encapsulated with Lin mLuc predominantly expressed Fluc protein in the liver (*30*). PL101 LNPs showed decreased liver expression, but increased spleen expression of Fluc protein, with a spleen-to-liver ratio of 0.42. Delivering Lin mLuc with PL40 LNPs further improved the preferential expression of Fluc protein in the spleen, with a spleen-to-liver ratio of 2.63. Interestingly, the encapsulation of Circ RNA favored spleen expression, with improved spleen Fluc protein expression and a two-fold increased spleen-to-liver ratio compared to the Lin counterpart (Fig. 3B). Furthermore, when delivered by PL40 LNPs, Circ mLuc presented prolonged expression in the spleen. Twenty-four hours post injection, 50% of mRNA expression was maintained in the spleen (Fig. 3C).

**Figure 3.**
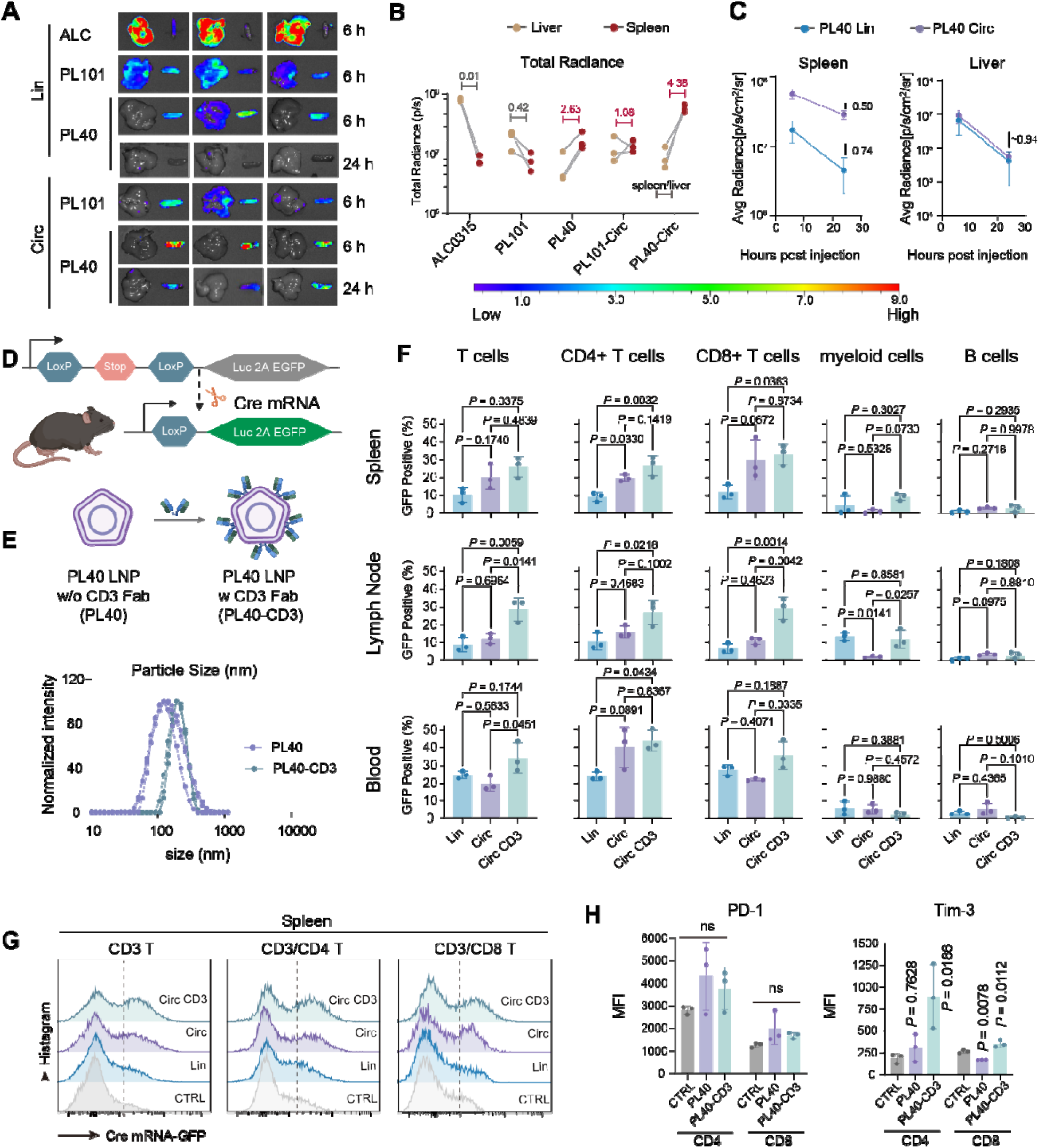
*In vivo* delivery of mRNA LNPs to immune cells. (**A**) IVIS images of *i.v.* injections of mLuc mRNA LNPs (mRNAs were prepared as Lin or Circ). 6 and 24 h after injection, organs were collected for imaging (n = 3). **(B)** Quantification of expression levels of mRNA LNPs 6 h after injection. **(C)** Comparison of expression levels of mRNA LNPs between 6 and 24 h. **(D)** Illustration of the mCre studies in loxP-Luciferase-2A-GFP mice. **(E)** LNPs were further modified with CD3-Fab and particle size were measured after modification (n = 3). **(F)** GFP expression in different immune cells within spleen, lymph node and blood were measured 48 h after *i.v.* injection of 20 µg mCre (Circ or Lin) in PL40 LNPs (with or without CD3-Fab modification), the background was subtracted (n = 3). **(G)** Representative histograms of T cells expression of GFP within spleens. **(H)** T cell gene expression markers after transfection with the mCre LNPs (n = 3). Data were represented as mean ± SD, “n” indicated biologically independent samples. Statistical significance was calculated through One-way ANOVA with Tukey test in F, and Dunnett test in H. All the schematic illustrations were created with BioRender.com.

To identify the specific cell types that expressed Fluc protein, we utilized a genetically engineered mouse model commonly used for *in vivo* mRNA delivery. This model carried a loxP stop codon cassette upstream of a GFP-2A luciferase sequence. Upon cleavage by cre-recombinase (delivered by mRNA encoding Cre, mCre), the mice could express both luciferase and GFP in the targeted cells (Fig. 3D). In this study, we compared PL40 LNPs delivered either Lin or Circ mCre. We also decorated PL40 LNPs with CD3-Fab using the maleimide NHS-thiol reaction, and set this as an antibody-targeted LNP control. The addition of the CD3-Fab resulted in a slight increase in particle size (approximately 20 nm), which is consistent with previous reports and confirms the successful conjugation of the antibody to the LNPs (*11*) (Fig. 3E). To analyze the cell types expressing the mRNA, cells from peripheral blood, lymph node and spleen were collected and analyzed by flow cytometry two days after *i.v.* injection of the mCre LNPs.

Interestingly, we found that both Lin and Circ mRNA LNPs, with (w) and without (w/o) CD3-Fab, primarily delivered and expressed mRNA in T cells, in all examined peripheral blood and peripheral lymphoid organs (Fig. 3F). Both non-targeted Circ mCre and Lin mCre showed comparable T cell expression of GFP in the blood and lymph node. However, the Circ mCre presented significantly higher (∼20%) transfection of spleen T cells. The addition of CD3-Fab increased T cell targeting, particularly in the lymph node (with ∼20% transfection), as compared to the non-targeted LNPs with ∼10% transfection. However, this advantage was not obvious in the spleen and blood as compared to the LNPs w/o CD3-Fab, confirming that PL40 LNPs could effectively deliver mRNA to T cells both *in vitro* and *in vivo* (Fig. 3F). We also observed that both CD4 and CD8 T cells took up the PL40 LNPs, w/ or w/o Fab decoration, at similar levels (Fig. 3G). It’s important to note that the addition of CD3-Fab has the potential to activate T cells and induce T cell tolerance (*31, 32*). To evaluate the state of T cells after LNP transfection, we examined the expression of Tim-3 and PD-1 markers. Though LNPs (w/ or w/o CD3-Fab) did not impact the expression of PD-1, a marker of the progenitor exhausted T cells, we observed a threefold increase in Tim-3 expression after dosing with CD3 Fab-decorated LNPs in CD4 T cells, suggesting a trend toward terminal exhaustion (Fig. 3H, fig. S5). To further reduce off-target delivery of LNPs to the liver, we also adjusted the formulation by replacing DOPE with DSPC or adding 1, 2-dioleoyl-sn-glycero-3-phospho-L-serine (DOPS) as the fifth component. However, these modifications led to a decreased T cell accumulation or overall mRNA expression *in vivo* (fig. S6). Therefore, the PL40 DOPE LNPs formulation was used for further investigations.

### PL40 LNPs efficiently engineered T cells into CAR Ts

Next, we aimed to reprogram T cells into CAR T cells for targeted cell lysis. We constructed four types of mRNA CAR cassettes (Fig. 4A-B): one encoding the canonical leukemia-specific 1928z CAR with the human CD3-CD28 intracellular domain (hCD19-hCAR) in a circular format, and the others encoding a mouse single-chain variable fragment (scFv) targeting mouse uPAR, along with either a human (muPAR-hCAR) in a circular format or mouse CD3-CD28 intracellular domain (muPAR-mCAR) in both linear and circular format. The hCD19-hCAR served as the control CAR in this study, as it is the most extensively investigated CAR-T cell product with nearly 30 ongoing clinical trials(*33, 34*).Both hCD19-hCAR and muPAR-hCAR delivered by PL40 LNPs exhibited expression levels of over 30% in human T cells (Fig. 4C and fig. S7). For the muPAR-mCAR mRNAs, we transfected them into mouse T cells. Compared to Lin mRNA with an 80% transfection efficiency, the Circ mRNA showed a 99% transfection efficiency 24 h after incubation (Fig. 4D). Notably, the Circ mRNA showed ∼10-fold higher mean intensity of the CAR protein compared to the Lin counterpart. Furthermore, the expression of the Circ mRNA also lasted longer, with around 15% of cells still expressing the CAR after 5 days. In contrast, the Lin mRNA dropped to <5% within 2 days (Fig. 4D). Subsequently, we evaluated the cytotoxicity of the LNP-transfected T cells. To demonstrate the targeting specificity for the antigens, we utilized both CD19 and uPAR as their respective control targets. By employing real-time *in vitro* cytotoxicity assays, we observed that both the hCD19-hCAR and muPAR-hCAR selectively lysed antigen-positive target cells (B cells for the hCD19-hCAR and 3T3-uPAR cells for the muPAR-hCAR) to around 90% or 60% respectively at an effector to target ratio (E:T) of 10:1, indicating that both CARs were functional as designed (Fig. 4E&F). We also tested the cytotoxicity of mouse CAR T cells transfected with muPAR-mCAR Circ RNA. We observed that the mouse CAR T cells lysed >95% 3T3-uPAR cells at an E;T ratio of 40 (Fig. 4G&H).

**Figure 4.**
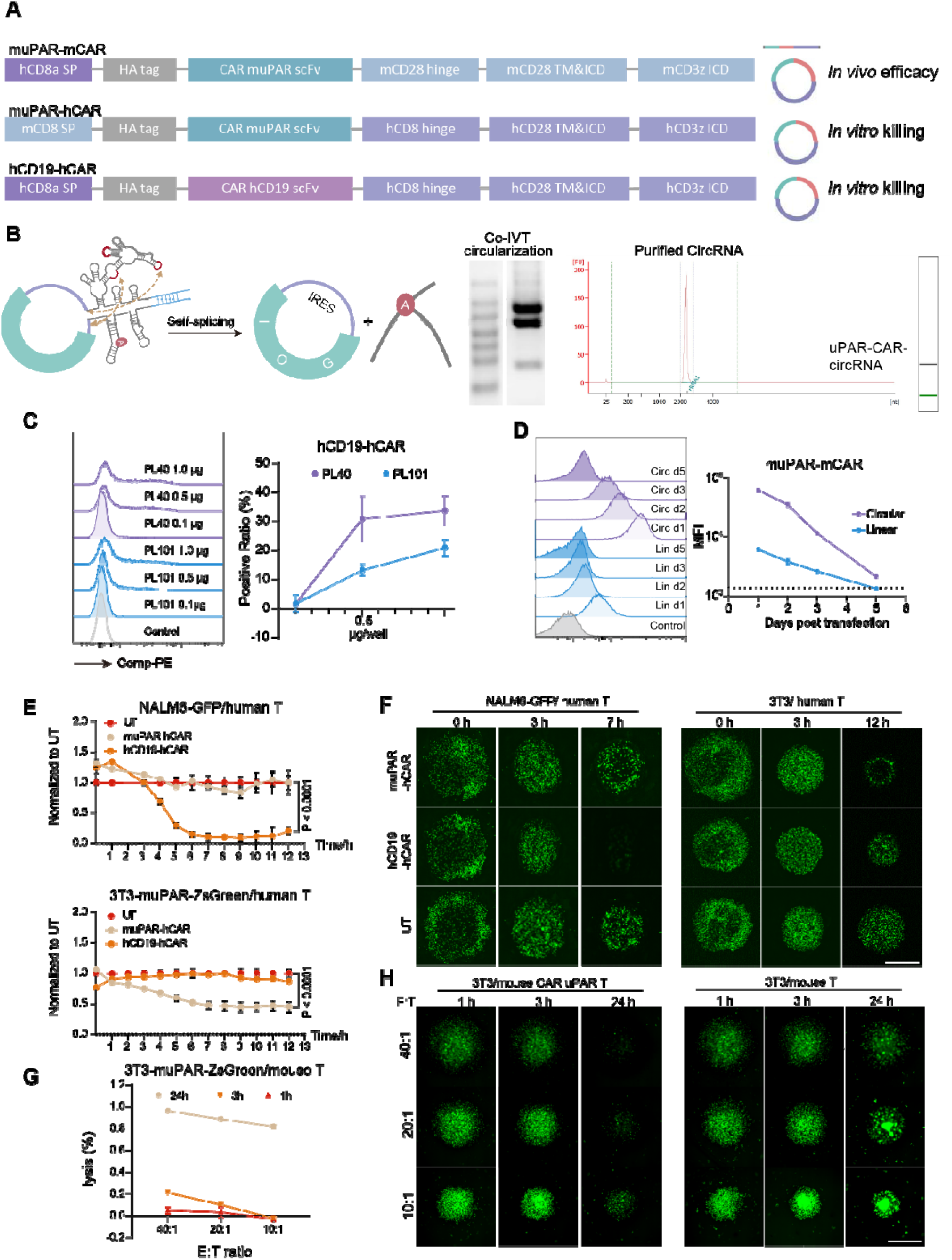
PL40 LNPs efficiently engineered T cells into CAR-Ts. **(A)** The construction map encoding CAR-m.uPAR m28z, muPAR-hCAR and hCD19-hCAR. The structure of mRNA (either Lin or Circ) were indicated in the figure. **(B)** The design, synthesis, purification and characterization of Circ RNA. **(C)** Transfection efficiency and dose dependence of CAMP LNPs encapsulating hCD19-hCAR mRNA. Human primary T cells were transfected with hCD19-hCAR mRNA at doses of 0.1 μg, 0.5 μg or 1 μg per million T cells that were encapsulated by PL101 or PL40 LNPs. The transfection efficiency was detected at 40 h using flow cytometry (n = 3, mean ± SD). **(D)** Expression of muPAR-mCAR mRNA in mouse T cells at different time points post transfection. Mouse T cells were transfected with Lin and Circ muPAR-mCAR mRNA that were encapsulated by PL40 LNPs at a dose of 0.5 μg/10^5^ T cells. Expression of the CAR were detected at 1, 2, 3 and 5 days by flow cytometry (n = 3). **(E)** The cytotoxicity of mRNA-based human CAR-T cells. The killing experiments were carried out at the E: T ratio of 10: 1. Representative images at different time points post coculture were presented in **(F)**, n=3 independent tests per group. **(E)** The cytotoxicity of mRNA-based mouse CAR-T cells. The killing experiments were carried out at the E: T ratio of 10: 1, 20:1 and 40:1. Representative images at different time points post coculture were presented in **(H),** n=3 independent tests per group. The whole process was monitored by IncuCyte SX5 (n = 3, scale bar = 1 mm), and the fluorescence intensity was calculated by IncuCyte. Data were represented as mean ± SD, “n” indicated biologically independent samples. Statistical significance was calculated through One-way ANOVA with Tukey test. All the schematic illustrations were created with BioRender.com.

### muPAR-mCAR PL40 LNPs show improved efficiency in liver fibrosis

Senescence contributes to a range of chronic tissue pathologies, including liver fibrosis where senescent HSCs contribute to the pathophysiology (*35*). Previous studies show that the *in vitro* prepared CAR-T cells targeting uPAR can treat liver fibrosis (*36*). Therefore, we adopted CCl_4_-induced liver fibrosis model to evaluate the *in vivo* therapeutic potency of the PL40 LNPs encoding CAR uPAR Cassette as proof of concept. Circ uPAR was used for *in vivo* efficacy study due to the superior expression (Fig. 4D). The C57BL/6J mice were treated with CCl_4_ for 5 weeks and then *i.v.* administrated with muPAR-mCAR Circ mRNA loaded in either CD3-Fab or non-Fab modified PL40 LNPs, using the blank vehicle and PBS as negative controls (Fig. 5A). After 4 doses of 30 µg/mouse mRNA delivered by PL40 LNPs w/o CD3-Fab, mice showed ∼ 2-fold reduced serum levels of alanine aminotransferase (ALT) compared to the PBS group, representing a reduction in liver damage (Fig. 5B). Moreover, the coverage of SA-β-gal and collagen in the liver both decreased by ∼2-fold (Fig, 5C), which indicated efficient elimination of pro-inflammatory senescent hepatic stellate cells (HSC). The multiplex immunohistochemical (mIHC) staining further showed that the levels of uPAR in livers from the non-Fab modified LNP-treated group was ∼4-fold lower, accompanied by a decrease in α-SMA (Fig. 5C). Mice treated with CD3-Fab modified LNPs presented a similar trend, but less reduction in collagen and uPAR^+^ cells (Fig. 5C&D). The Th-1 cytokines IFNγ and IL-2 expression by T cells in the blood of LNP-treated mice was approximately 1.5 times higher than the PBS group (Fig. 5E), suggesting a trend of T cell activation. Furthermore, T cell exhaustion markers such as PD-1 and Tim3 were obviously increased after treatment with CD3-Fab LNPs, which were not upregulated significantly in the non-Fab group (Fig. 5F). This is consistent with the Loxp-GFP study (Fig. 3H), where we found adding CD3-Fab increased T cell exhaustion.

**Figure 5.**
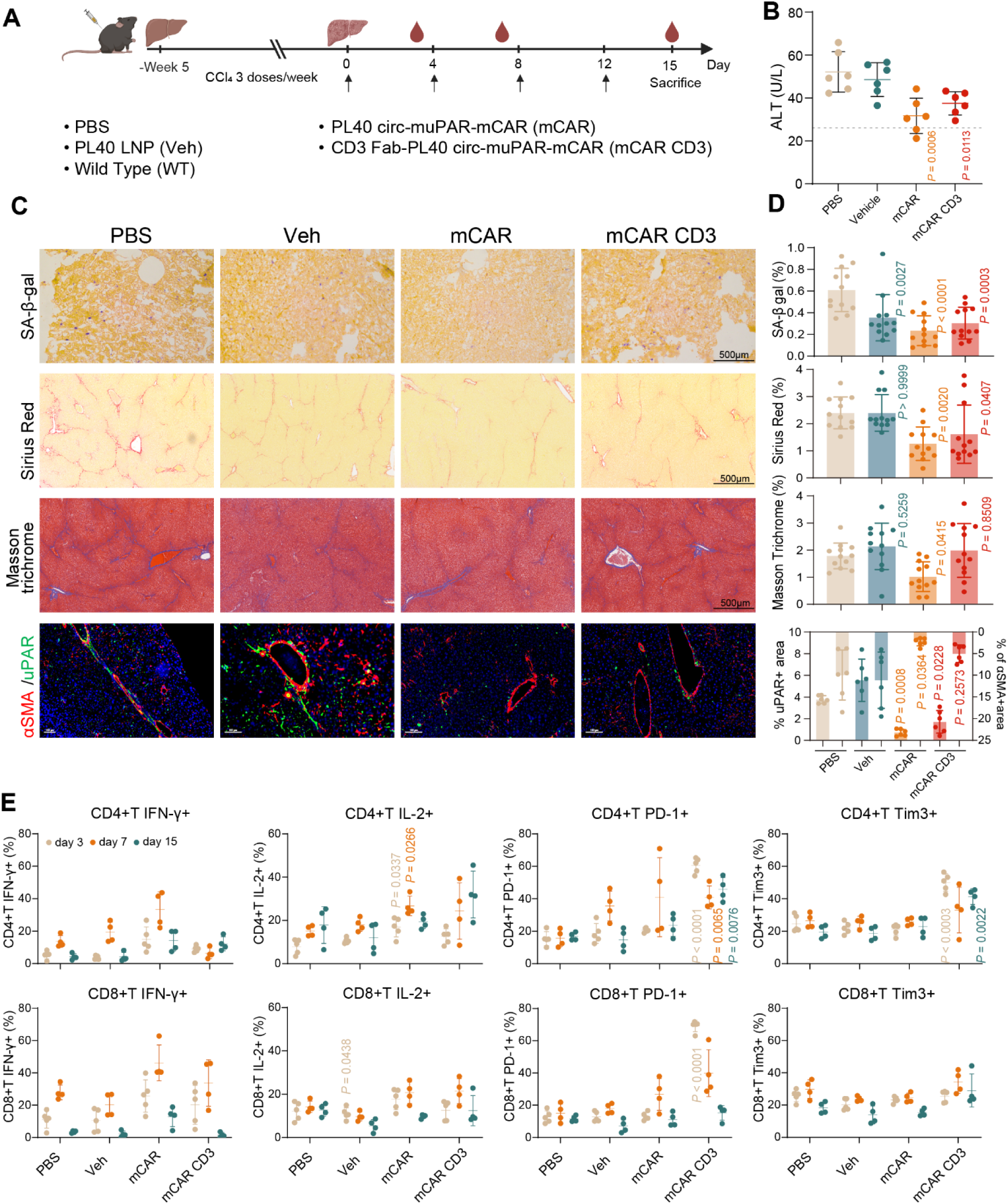
Anti-fibrosis effects of muPAR-mCAR LNPs in CCl_4_-induced fibrosis. **(A)** Schematic representations of the CCl_4_-induced liver fibrosis models and treatment schedules. muPAR-mCAR encapsulated by CD3-Fab or non-Fab modified PL40 LNPs were *i.v.* administrated to mice at a dose of 30 µg/mouse. **(B)** Serum ALT from CCl_4_-induced fibrotic mice with all groups after 4 doses treatment (n = 6, mean ± SD). **(C)** Representative mIHC and fluorescent stainings of SA-β-gal, Sirius red, Masson’s trichrome, and uPAR/αSMA in the livers of CCl_4_-induced liver fibrotic mice with all groups after treatment. The quantifications of the coverage% area of SA-β-gal, collagen and uPAR/αSMA were performed in 2 randomly selected fields per mouse (from n = 6 biological independent mice per group, mean ± SD). **(D)** IFNγ, IL-2, PD-1 and Tim3 expression of T cells in the blood of treated mice measured by flow cytometry at day 4, day 7 and day 15 (n = 4 per time points, mean ± SD). n” indicated biologically independent samples. Statistical significance was calculated through One-way ANOVA with Dunnet test. All the schematic illustrations were created with BioRender.com.

Based on these results, the non-Fab LNPs presented superior anti-liver fibrosis efficacy compared with the CD3-Fab modified LNPs, thus the subsequent validation focused on non-Fab modified LNPs. Further analysis showed that the infiltration of overall T cells, especially the effector memory T cells, increased markedly in non-Fab LNPs CAR group compared to PBS group (3.2 folds for CD8^+^ T cells and 8.1 folds for CD4^+^ T cells) (Fig. 6A). An increase in M2 macrophage infiltration was also observed, revealing an anti-inflammatory microenvironment persisted after Car T-treatment.

**Figure 6.**
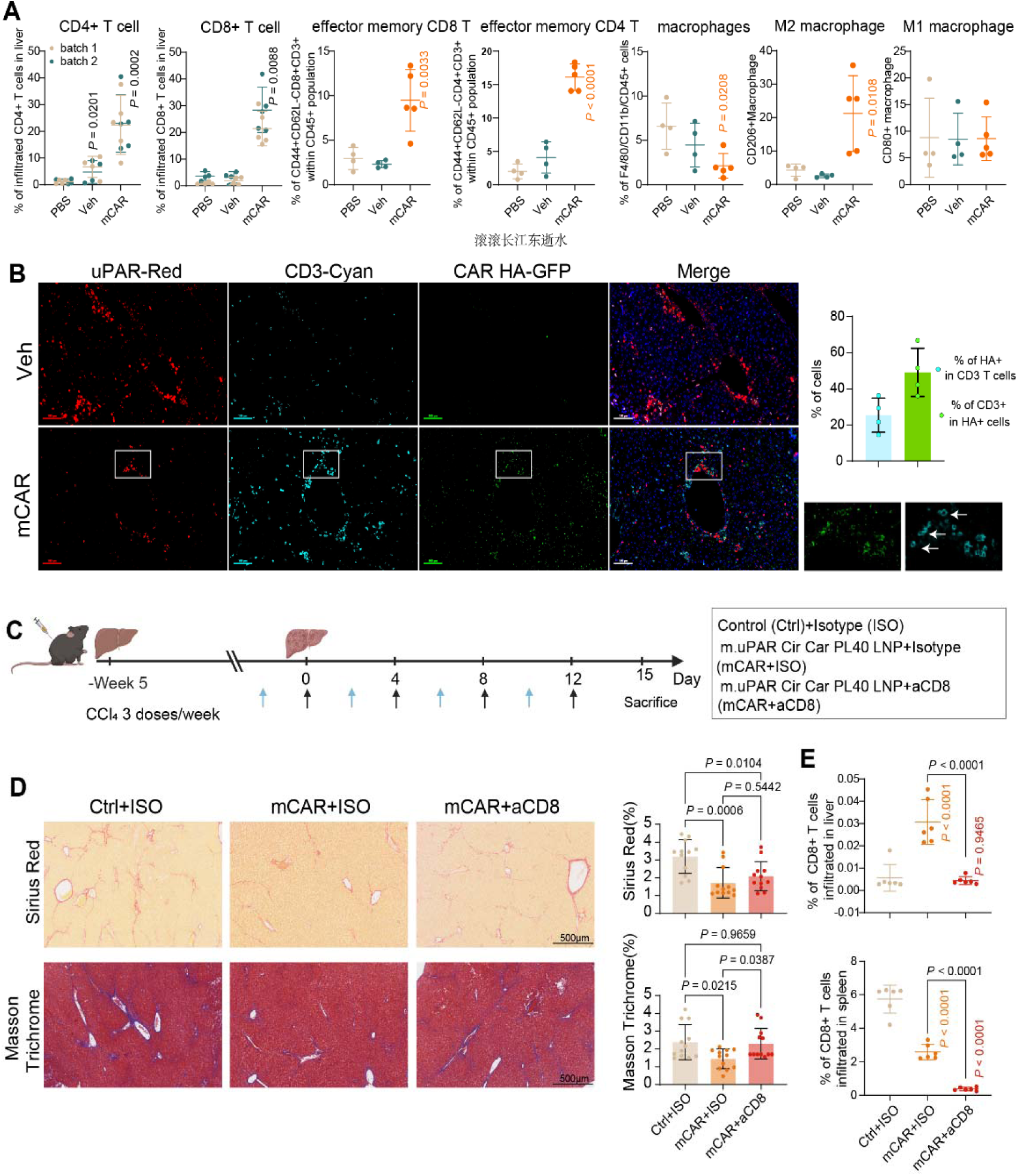
Evaluation of the function of Car T cells in liver fibrosis treatment. **(A)** Infiltrated T cells and macrophages proportion in the liver of treated mice in Fig. 5A measured by flow cytometry at day 15. n = 4 mice in PBS and Veh group, n = 5 mice in mCAR group, mean ± SD. Experiments were repeated in two batches with similar results. **(B)** Multi-color fluorescent staining of uPAR (red), CD3 (cyan) and HA-tag (green) in the livers of treated mice, n = 4 mice per group. Quantifications were shown on the right. **(C)** Schematic representations of the CCl_4_-induced liver fibrosis models and treatment schedules. aCD8 was i.v. administrated to deplete the endogenous T cells. muPAR-mCAR encapsulated by non-Fab modified PL40 LNPs were *i.v.* administrated to mice at a dose of 30 μg/mouse (n = 6 mice). **(D)** Representative IHC staining of SA-β-gal, Sirius red and Masson’s trichrome in the livers of treated mice with all groups in Fig. 6C after treatment. The quantification of the coverage% area of SA-β-gal and collagen was performed in 3 randomly selected fields per mouse (from n=6 mice per group, mean ± SD). **(E)** Infiltrated CD8^+^ T cells in the liver of treated mice in Fig. 6C measured by flow cytometry at day 15 (n = 6 mice). “n” indicated biologically independent samples. Statistical significance was calculated through One-way ANOVA with Tukey test. All the schematic illustrations were created with BioRender.com.

To confirm that the therapeutic effect was induced by the killing of uPAR-expressing senescent cells by the *in vivo* engineered CAR-T cells, the colocalization of HA-tagged Car T with uPAR^+^ cells were visualized through mIHC staining. Compared with the LNPs vehicle group, the HA^+^ CAR T cells were infiltrated into liver parenchyma, neighboring uPAR^+^ cells in the CAR PL40 LNPs group (Fig. 6B). Furthermore, depletion of the endogenous T cells with an anti-CD8 antibody (Fig. 6, C&E), abolished the efficacy of the LNPs treatment (Fig. 6D). Collectively, these results demonstrated efficient treatment of liver fibrosis by non-Fab modified CAR uPAR LNPs.

### muPAR-mCAR PL40 LNPs shows improved efficiency in rheumatoid arthritis *in vivo*

Besides fibrosis, uPAR had been witnessed to possess multifactorial approach in mediating Rheumatoid Arthritis (RA) pathogenesis, in which uPA secreted by neutrophils, chondrocytes, and monocytes interact with uPAR (*37–40*). To further investigate the localization of uPAR in different subcellular populations and its correlation between senescent phenotypes, we integrated and compared relevant Bulk RNA-Seq and scRNA-Seq data of synovial membrane of RA patients from public databases(*33, 34*). Both data shows that uPAR were highly expressed in fibroblast and monocytes (Fig. 7A, fig. S8&9). Further, Bulk RNA-Seq and scRNA-Seq data shows a relatively clear co-localization of uPAR with senescence makers such as GLB1 and SPERPINE1 in stromal cells and myeloid cells, as well as several Senescence-Associated Secretory Phenotype (SASP) at the subcellular levels (Fig. 7A, Fig. S8&S9). A collagen-induced arthritis (CIA) mouse model based on DBA/1 mice was further established as previously reported to confirm the efficacy of our in-vivo CAR-T therapy targeting uPAR-expressing senescent and inflammatory cells in RA process (Fig. 7B) (*41, 42*). Consistent with clinical samples, we observed that uPAR was mainly expressed in synovial myeloid cells and partially expressed in spindle cells (likely to be stroma cells) (Fig. 7H). We adopted *i.p.* injection of MTX which is widely used in RA treatment at a dose of 5 mg/kg as a positive control. The severity of arthritis was evaluated in terms of the changes in the main symptoms of CIA like swelling and redness. During the treatment schematic, the clinical scores and hind paw thickness of the mCAR PL40 LNPs treated group decreased after each administration and the scores were ultimately ∼2.4 times lower than control mice or mice receiving blank PL40 LNPs vehicle (Fig. 7C to E), manifesting a reduction in inflammation and edema. H&E stained sections showed that the level of inflammation decreased, and the joint cavity was restored, with clear interfaces, no obvious synovitis, articular cartilage degeneration in the ankles and toxicity in organs of muPAR-mCAR PL40 LNPs treated mice (Fig. 7F, fig. S10). On the contrary, severe pathological changes, including extensive inflammatory cell infiltration, synovial hyperplasia and cartilage erosion, were observed in the control group. Additionally, safarin O/Fast green staining showed optimal preservation of the cartilage structure of the mice in the mCAR PL40 LNPs –treated group (Fig. 7F). The mIHC analysis indicated a significant decrease of uPAR coverage and an increase of infiltrated T cells neighboring uPAR^+^ cells in the mCAR PL40 LNPs group, demonstrating that the CAR induced T cell-mediated clearance of senescent cells. Consistently, the infiltration of anti-inflammatory M2 macrophage also increased by ∼6-folds at the same time (Fig. 7H&I).

**Figure 7.**
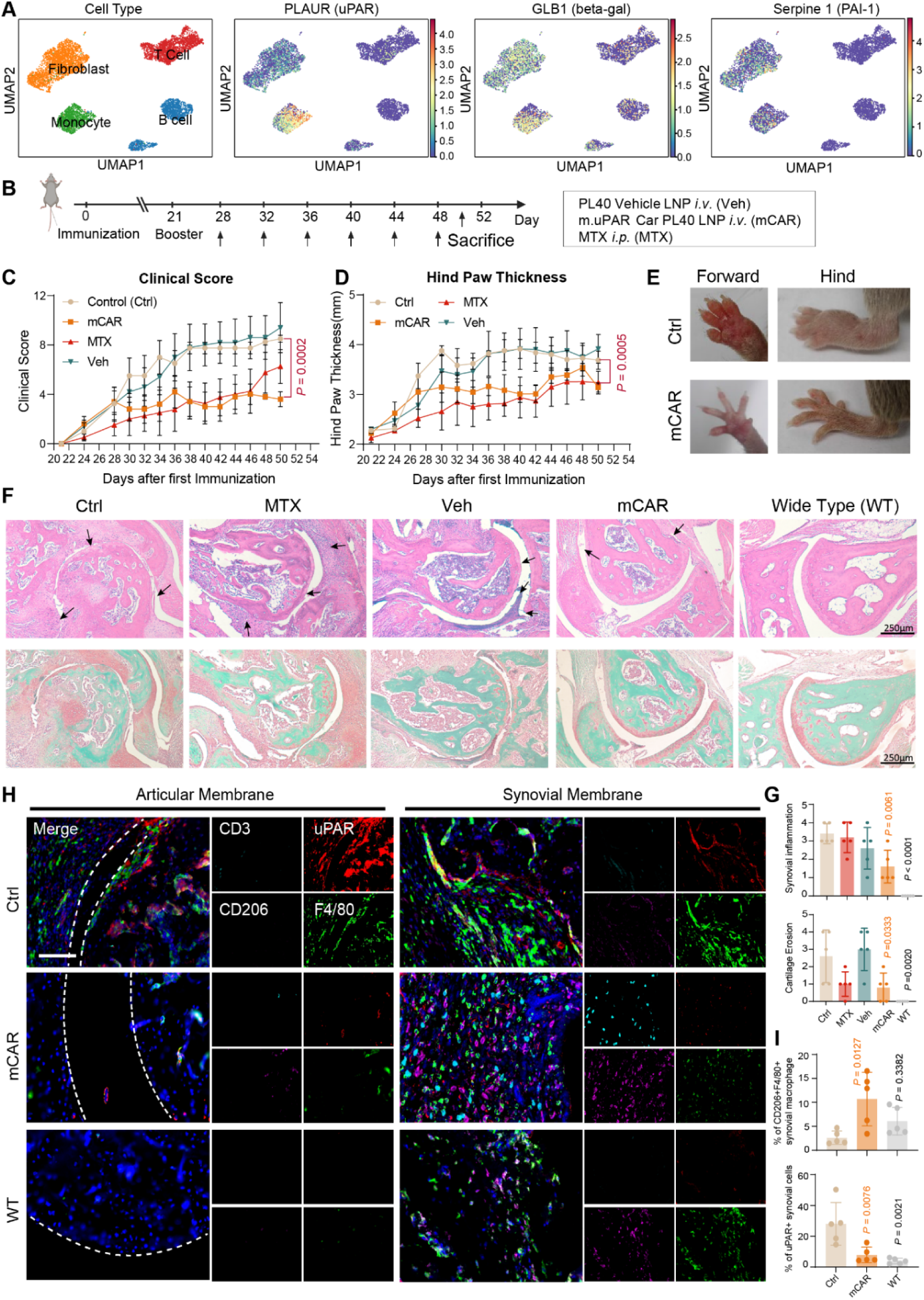
*In vivo* therapeutic effect of muPAR-mCAR LNPs in collagen-induced arthritis. **(A)** scRNA-Seq data of synovial membrane from RA patients from public resources. **(B)** Schematic diagram of the treatment schedule for the CIA mouse model. **(C** to **E)** Clinical score and paw thickness of the CIA mice in different treated groups (n = 5, mean ± SD). **(F)** Representative of H&E and safarin O/Fast green staining in the ankles of CIA mice with all groups. The quantification of inflammation level and cartilage erosion was performed in each mouse (from n = 5 biological independent mice per group, mean ± SD) and shown in **(H)**. **(G)** Paraffin sections mIHC staining of uPAR (red), CD3 (green), F4/80 (blue) and CD206 (purple) in the ankles of CIA mice. Scale bar 100 µm. Quantifications of the mIHC stainings were presented in **(I)**, from n = 5 individual samples. “n” indicated biologically independent samples. Statistical significance was calculated through One-way ANOVA with Dunnett test. All the schematic illustrations were created with BioRender.com.

These results collectively suggested that the CAR-muPAR PL40 LNPs therapy could edit and mobilize T cells to kill senescent cells displaying uPAR in the joint of CIA mouse without causing inflammation, thereby effectively reestablish joint immune homeostasis.

### Screening and optimization of humanized CAR uPAR mRNA for clinically relevant applications

We further screened two mouse monoclonal antibodies against human uPAR with high affinities from more than 60,000 mouse hybridoma clones (Fig. 8A&B, fig. S11A to C). AB4 and AB20 bound to the epitope in D2 domain and conformational epitope in D2-D3 domains of uPAR, respectively, while the commercial uPAR antibody Vim5 bound to the D1 domain (Fig. 8C&D). These allowed AB4 and AB20 to recognize either intact or cleaved uPAR when it was activated by uPA (Fig. 8E&F). To avoid immunogenicity, the sequences of both heavy and light chains of AB4 and AB20 antibodies were humanized and transfered to scFv format for CAR (Fig. 8F). The combination VH2+VL1 CAR of AB4 (CAR#4) and the VH1+VL2 of AB20 (CAR#20) have been selected based on their binding affinities to uPAR (Fig. 8G). The difference of the binding affinity between these two scFvs was similar to that of their full-length antibodies (Fig. 8H&D). Next, we evaluated the cytotoxicity of lentivirus-delivered CAR#4 and CAR#20 T cells by targeting uPAR^+^ AGS and THP-1 cells. We found that both CAR#4 and CAR#20 were potent in killing uPAR^+^ target cells but not uPAR-deficient cells (Fig. 8I&J, fig. S11D). Moreover, to compare the cytotoxicity of transit-expressed CAR by LNP to chromosome-integrated CAR by lentivirus, we constructed Lin and Circ RNA of CAR#20, and found that all of three types of CAR had high potency and efficacy in killing uPAR^+^ target cells but not uPAR-deficient cells (Fig. 8K). Interestingly, CircRNA-based CAR#20 showed better cytotoxicity than both LinRNA-based and lentivirus-based CAR#20. Together, these results have shown that RNA-based anti-human uPAR CAR T cells were also potent and specific in the elimination of uPAR^+^ target cells, which provides the potential for further clinical translation.

**Figure. 8.**
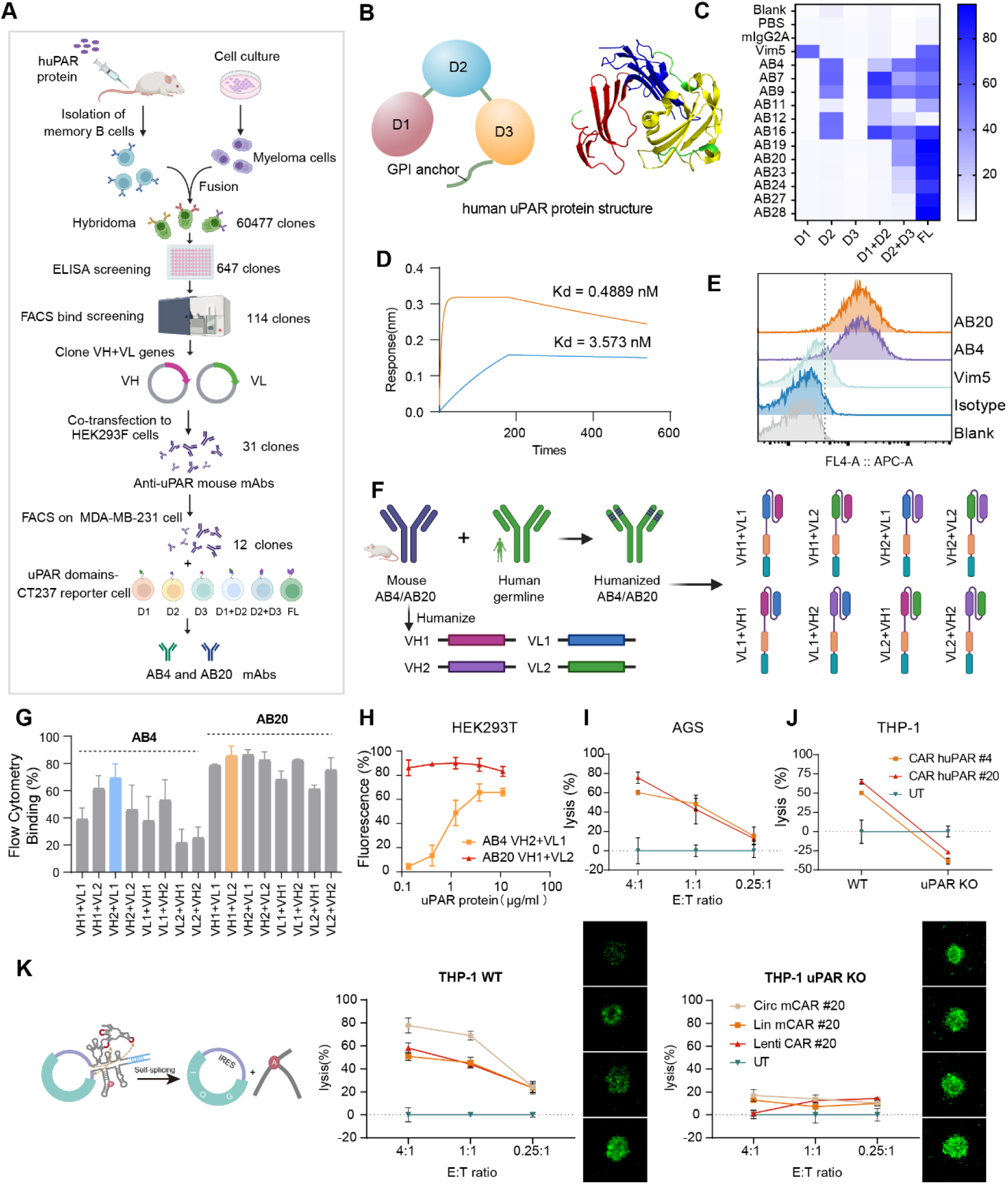
Preparation of uPAR monoclonal antibodies and construction of uPAR CAR-T cells. **(A)** Schematic of the strategy for preparing uPAR mouse monoclonal antibodies. **(B)** The structure of the human uPAR protein. The human uPAR has 3 domains and a GPI anchor. **(C)** Screen mAbs against different fragments of human uPAR. Flow cytometry was used to detect the activation of CT237 reporter cells with different domains of chimeric uPAR extracellular region (D1, D2, D3, D1+D2, D2+D3 and FL) **(D)** The binding affinity of AB4 and AB20 monoclonal antibodies to human uPAR measured by SPR. **(E)** Compared with the commercial uPAR antibody Vim5, AB4 and AB20 mAbs more sensitively detects uPAR on THP-1 cells. **(F)** Schematic illustrates the humanization of AB4 and AB20 monoclonal antibody, and construction of the fully humanized CAR cargo. **(G)** HEK 293 cell display of humanized anti-uPAR scFV constructs and the binding affinity tests with target human uPAR antigen. Screening identified 2 humanized constructs with higher binding capability (n = 3, mean ± SD). **(H)** The dose dependent binding properties of humanized AB4 and AB20 monoclonal antibodies to uPAR protein measured by flow cytometry (n = 3, mean ± SD). **(I)** The cytotoxicity of lenti-virus transfected AB4 VH2+VL1 CAR-T cells and AB20 VH1+VL2 CAR-T cells against upar^+^ AGS cell line at different E: T ratio (n = 3, mean ± SD). **(J)** The cytotoxicity of lenti-virus transfected AB4 VH2+VL1 CAR-T and AB20 VH1+VL2 CAR-T cells were tested against wild-type and uPAR-knocked out THP-1 cell lines, at E: T ratio of 4: 1 (n = 3, mean ± SD). **(K)** Circ and Lin RNA of AB20 CAR were designed. The cytotoxicity of AB20 VH1+VL2 CAR T cells, transfected by Lin RNA, Circ RNA PL40 LNPs, or lenti-virus were tested and compared on wild-type and uPAR-knocked out THP-1 cell lines, at different E: T ratio (n = 3, mean ± SD). Statistical significance was calculated through One-way ANOVA with Tukey test. All the schematic illustrations were created with BioRender.com.

## DISCUSSION

CAR-T therapy has achieved a remarkable clinical efficacy to hematopoietic malignancies. However, conventional CAR-T therapy requires a highly complex and personalized engineering process to produce CAR T cells *ex vivo and* carries potential risks of T cell malignancies or autoimmune disorders. The mRNA-based *in vivo* CAR T cell therapy may overcome these limitations.

In this study, we identified a specific lipid, PL40, that belongs to a new class of CAMP ionizable lipids and can efficiently deliver mRNA to T cells when formulated into an LNP platform, without requiring antibody modifications against T cells. Physical characterization revealed the CAMP LNPs exhibit an irregular polygonal morphology and high rigidity. Particle shape is known to influence endocytosis, with macrophages preferring spherical particles over those with higher aspect ratios or irregular geometries (*43–45*). Interestingly, our results show T cells exhibit the opposite preference, demonstrating enhanced uptake of rigid, irregularly shaped LNPs. The multi-faceted surface morphology of PL40 LNPs likely arises from phase separation between lipid domains. This structural disorder may facilitate membrane fusion and interaction with phase-separated lipid rafts in the T cell membrane, further promoting endocytosis (*46*).

Consistent with this, the PL40 LNPs exhibited higher cellular uptake and mRNA expression in T cells compared to more uniform conventional LNPs. Previous studies have also shown T cells prefer rigid objects, which can enhance activation and metabolism, enabling greater cellular uptake (*47–51*).

While the non-antibody modified PL40 LNPs have demonstrated comparable or better efficacy than CD3 Fab-targeted formulations, their targeting specificity may be less precise. However, the ability to simultaneously transfect other immune cell types could lead to synergistic improvements in T cell function. Further exploration of the mechanisms underneath the *in vivo* T cell specific accumulation of CAMP lipids and developing multi-modal targeting strategies may enhance the therapeutic potential of LNP-mediated mRNA delivery to T cells.

To further prolong mRNA expression *in vivo*, we introduced CircRNA. Previous studies have shown that circularization of mRNA prevents enzymatic degradation, leading to prolonged expression. Consistently, CircRNA prolonged mRNA expression in T cells and the spleen in our study, particularly improving the spleen-to-liver ratio of protein expression. Whitehead *et al.* have shown that the expression of base-modified mRNAs, including m1ψ and m5C/ψ, was enhanced in the spleen but not significantly changed in the liver (*52*). This result is consistent with the observed effects of CircRNA, suggesting immune cells may be more sensitive to changes in RNA structure. The underlying mechanism for the preferential expression of CircRNA in the spleen still needs further investigation.

Emerging evidence has shown that cellular senescence is associated with chronic inflammatory diseases, such as liver fibrosis and rheumatoid arthritis (RA) (*53, 54*). Clinical trials have focused on chemical senolytic agents, such as the combination of Dasatinib and Quercetin, which eliminate senescent cells by intervening in their apoptotic environment (*55, 56*). However, these senolytic drugs have potential off-target toxicity (*57*), and improving their specificity will be crucial. Antibody-equipped cell therapy provides a strategy with high specificity to remove senescent cells. uPAR has been recently identified as a surface marker of senescent cells in liver fibrosis and lung cancer (*4*). Our results have shown uPAR is expressed on both senescent fibroblasts and monocytes in human RA patient samples and a mouse RA model, in addition to liver fibrosis. The non-antibody modified CAMP lipid-engineered the CAR-uPAR T cells *in vivo* efficiently eliminated senescent cells and alleviated disease symptoms. Further clinical investigation is emergently in need of the promising *in vivo* senolytic CAR-T therapy. In summary, our data highlights the potential of using a non-antibody targeting strategy to engineer T cells *in vivo* for senolytic therapy of inflamm-aging related diseases.

## MATERIALS AND METHODS

### Sudy design

This study aimed to innovate lipid-based mRNA delivery by developing a new class capable of selectively transfecting T cells in vivo without relying on antibody modification. The objectives were multifaceted: (1) to design and screen a lipid library for precise RNA delivery to primary human T cells in vitro, examining the specificity of this interaction; (2) to apply the most effective lipid, PL101, for in vivo delivery of linear and circular RNA, assessing its distribution and the benefits of circular RNA at the organ and cellular levels; (3) to utilize PL101 LNPs for delivering muPAR-mCAR mRNA as a therapeutic approach for liver fibrosis and rheumatoid arthritis; and (4) to screen and validate the functionality of humanized CAR against human uPAR, advancing its potential for clinical use. Experimental group sizes were calculated based on prior research, and all pathology analyses were conducted by a researcher blinded to the study conditions. Methodological details, including replicate counts, statistical methods, and P values, are delineated in the figure legends.

### Materials

1,4-Bis(3-aminopropyl)piperazine, undecan-1-ol, (9Z,12Z)-hexadeca-9,12-dien-1-ol, 3-allyloxy-1,2-propanediol, 1,8-octanediol, 4-methoxybenzylchloride (PMBCl), undecan-3-ol, 2,3-dichloro-5,6-dicyano-1,4-benzoquinone (DDQ), heptadecan-9-ol were purchased from J&K Scientific (China). 3-Bromopropylamine hydrobromide, ethanolamine, 3-aminopropan-1-ol, 3-(4-methylpiperazin-1-yl)propan-1-amine, 4-(2-aminoethyl)morpholine, 3-phenylpropan-1-amine, 2-methyl-3-(pyrrolidin-1-yl)propan-1-amine, 3-dipropylaminopropylamine, 2-(piperazin-1-yl)ethan-1-amine, N,N-bis(3-aminopropyl) methylamine were purchased from Bide Pharmatech Co., Ltd. Tetrakis (triphenylphosphine) palladium, barbituric acid, N,N’-dicyclohexylcarbodiimide (DCC), 4-dimethylaminopyridine (DMAP), sodium hydride (NaH), phosphorus oxychloride were purchased from Innochem (China).

All the ionizable lipids including DLin-MC3-DMA (MC3) and 6-((2-hexyldecanoyl)oxy)-N-(6-((2-hexyldecanoyl)oxy)hexyl)-N-(4-hydroxybutyl)hexan-1-aminium (ALC-0315)were purchased from Avanti Polar Lipids, Inc. Helper lipids including cholesterol, 1,2-Dioctadecanoyl-sn-glycero-3-phophocholine (DSPC), 1,2-Dioleoyl-sn-glycero-3-phosphoethanolamine (DOPE), 1,2-dimyristoyl-rac-glycero-3-methoxypolyethylene glycol-2000 (DMG-PEG2000) and 1, 2-dioleoyl-sn-glycero-3-phospho-L-serine (DOPS) were bought from A.V.T. Pharmaceutical Tech Co., Ltd. Circular mRNA (mLuc) was provided by Circode. All the plasmid sequences were provided by Genscript Co.,Ltd. Cell lines were purchased from Procell Life Science & Technology Co.,Ltd. Fetal bovine serum was purchased from Thermo Fisher Scientific Co.,Ltd. Other reagents for basal culture were bought from Meilunbio Co., Ltd. The human PBMC was purchased from Milestone® Biotechnologies.

## Methods

### CAMP lipid synthesis

The synthesis of the CAMP lipids was carried out using POCl_3_ as the starting compound. Three nucleophilic reagents (two lipid alcohols and one amine) were added sequentially in solvent CH_2_Cl_2_ with the presence of triethylamine. The detailed procedure is as follows: First, two lipid alcohols and one amine were separately dissolved in CH_2_Cl_2_, and triethylamine (1 equivalent) was added to each solution as a preparation step. Next, the solution of the first lipid alcohol was added to a solution of POCl_3_ under 4 °C and the reaction was allowed to proceed for 3 hours.

Then, the solution of the second lipid alcohol was added to the reaction mixture, and the reaction was continued at 4 °C to room temperature for 2 hours. Finally, the solution of the amine was added, and the reaction was further continued at room temperature for 2 hours. After the completion of the reaction, the excess amine and the generated byproduct, the triethylamine salt, were removed by washing with saturated sodium chloride solution. The product was then purified by column chromatography.

Using the above method, the lipids PL15, PL16, PL101, PL102, PL39, PL40, PL48, PL66, and PL68 were synthesized. For the synthesis of lipids PL49, PL50, PL51, PL52, PL63, PL64, PL65, PL67, PL69, and PL70, the same method was used, but different amino head groups were selected for further bromination.

The structure of the synthesized lipids was confirmed by ^1^H NMR spectrometry, ^13^C NMR spectrometry (Bruker AVANCE-400 NMR spectrometer with a Magnex Scientific superconducting magnet) and mass spectrometry (Waters Xevo G2 QTOF and ThermoFisher Orbitrap Exploris).

### Linear mRNA synthesis and purification

Lin mRNAs encoding CAR-m.uPAR m28z, GFP and firefly luciferase were synthesized by *in vitro* transcription (IVT). A linearized pUC57 plasmid vector, which contained a T7 promoter, 5’ untranslated region (5’UTR), a coding sequence (CDS) encoding each mRNA mentioned above, 3’UTR and a poly A region (∼100nt), were used as a template for transcription. Clean-cap AG 5’ capping (Cap 1), and 100% 1-methylpsuedo-uridine UTP were used during IVT to improve protein translation efficiency and minimize immunogenicity. The IVT reactions were performed according to the manufacturer’s instructions (Hongene Biotech Inc., China). Then, the mRNA was purified by MagicPure RNA Beads (TransGen Biotech, #EC501).

### Circular mRNA synthesis and purification

The coding sequences of each protein were inserted into backbone plasmid to construct Circ RNA vectors. RNAs were IVT from the XbaI (Thermo, FD0685)-digested linearized plasmid DNA template using a MEGAscript™ T7 (Thermo, AMB1335-5) in the presence of unmodified NTPs. After DNase I treatment, the IVT products were column purified with an RNA Clean and Concentrator Kit (ZYMO Research, R1019) to remove excess NTP and other salts in IVT buffer.

The total RNA products were subjected to a series of purification steps to gain high purity of Circ RNA. First, affinity chromatography was conducted using a HiTrap NHS-activated HP column (Cytiva, 17071601) coupled with a specific ligand to selectively remove precursor and intron RNAs. The binding buffer contained 15mM LiCl, 10mM Tris, 0.5mM EDTA, and water for injection. Second, the RNA underwent Fast Protein Liquid Chromatography (FPLC) employing size exclusion chromatography (Sepax, 215950-30030) to remove polymer and small RNA at a flow rate of 10 ml/min. The elution buffer was composed of 75 mM phosphate buffer (PB), 10 mM Tris, 0.5 mM EDTA, and water for injection, pH adjusted to 7.4. Finally, optional RNase R treatment was performed to remove nick RNAs. The reaction conditions were 37°C for 30 min. The purified CircRNAs were subjected to further experiments after column purification with an RNA Clean and Concentrator Kit (ZYMO Research, R1019). The integrity of the Circ RNAs were confirmed using the Agilent 2100 Bioanalyzer (Agilent Technologies).

### CAMP lipid nanoparticle formulation and characterization

LNPs were prepared either by hand mixing or microfluidic mixing as previously described. Briefly, an aqueous solution of the mRNA and an ethanolic solution of the lipid components were mixed at a ratio of 3:1, respectively. The ethanol phase consists of cationic lipid (CAMP lipid), 1,2-distearoyl-sn-glycero-3-phosphocholine (DSPC, AVT, China) or 1,2-dioleoyl-sn-glycero-3-phosphoethanolamine (DOPE, Avanti, USA), cholesterol (AVT, China) and 1,2-dimyristoyl-sn-glycerol, methoxy polyethylene glycol (DMG-PEG2000, AVT, China). The aqueous phase was prepared in 10 mM citrate buffer (pH 4). After mixing, the obtained LNPs with a final mRNA concentration of 0.1 mg/mL were dialyzed in PBS in a dialysis bag (12-14 kDa) at 4 °C overnight. The mRNA concentration and encapsulation efficiency of the LNPs were measured using the Quant-iT RiboGreen RNA assay (Invitrogen). The hydrodynamic diameter and zeta potential were measured by dynamic light scattering (Zetasizer Nano ZSP, Malvern), with the samples diluted in 1× PBS or 4 mM KCl, respectively. To evaluate the plasma stability of the LNPs, they were incubated in fetal bovine serum (FBS) for 6 hours at 37 °C with 200 rpm agitation. The particle sizes of the LNPs were measured by a Zetasizer at 0 h, 1 h, and 6 h.

### Atom Force Microscopy (AFM)

AFM imaging and force measurements were carried out in PBS at room temperature by using Bruker MultiMode 8, PeakForce QNM in Fluid mode. The LNP was incubated on a mica plate at room temperature for at least 20 min in a humidified box. The surface of mica plate was gently washed one time with PBS, and 20 µL PBS was dropped to the mica surface before measurement. The probe used for measurement was SCANASYST-FLUID+(Bruker), and before measurement, 40 μL PBS was used to wet the probe. The data was analyzed by NanoScope Analysis 1.9 software.

### Small Angle X-ray Scattering (SAXS)

Small-angle X-ray scattering (SAXS) experiments at 20 °C were conducted using a Xeuss 2.0 SAXS system (Xenocs SA) with a two-pinholes collimation setup for the LNPs at various concentrations. X-ray radiation with a wavelength of 1.5418 Å was generated using a Cu Kα X-ray source (GeniX3D Cu ULD). The sample-to-detector distance is ca. 600 mm, corresponding to a q-range of 0.16 to 8.19 nm^-1^. The beam diameter was approximately 1 mm. The LNPs, which were first condensed to a concentration of 0.4 mg/mL, were loaded into quartz capillary tubes (WJM-Glas/Müller GmbH) with a 2 mm diameter, 80 mm length, and 0.01 mm wall thickness, and sealed at the top with epoxy resin. The 2D SAXS patterns were collected for an image acquisition time of 1800 s under vacuum using a semiconductor detector (Pilatus 300K, DECTRIS) with a resolution of 487×619 pixels (pixel size of 172×172 μm^2^). The obtained SAXS patterns were corrected for detector noise, air scattering, and sample/quartz-capillary absorption. 1D intensity profiles were then integrated from the background-corrected 2D SAXS patterns. The long-distance (L) was determined using the q-value at the first intensity maximum (denoted as q_m_) in the 1D SAXS intensity profiles, according to the formula: L = 2π/q_m_.

### Cryo-transmission electron microscopy (cryo-TEM) measurement

The morphology of LNPs was characterized by cryo-electron microscopy (cryo-TEM). For cryo-TEM measurement, the LNPs were dialyzed in 20 mM Tris (pH 7.4) containing 8% sucrose at 4°C overnight, and then condensed to 0.4 mg/ml by ultrafiltration. Cryo-TEM image was acquired using Themis 300 (Thermo Fisher Scientific).

### *In vitro* Luciferase mRNA transfection

The human primary T cells were isolated from human PBMCs and plated in white, clear-bottom 96-well plates at a density of 1×10^5^ cells per well. Three days after activation, LNPs containing mLuc RNA were added to cells at 60 ng mRNA per well. After 24 h transfection, the transfection efficiency was measured by Firefly-Glo Luciferase Reporter Assay Kit (Yeasen Biotechnology Co., Ltd.) according to the manufacturer’s protocol, using Biotek synergy H1 microplate reader. To transfect mouse T cells, activated primary mouse T cells or CT237 were resuspended at a density of 1×10^5^ cells/m. Then, LNPs containing CAR RNA were added to cells at 5 μg mRNA per million cells. The CAR expression was detected by flow cytometry after 24 h of incubation (*58*).

### *In vitro* cell uptake

Primary human T cells were plated in 96-well plates at a density of 10^5^ cells per well. The cells were pre-incubated with small molecule endocytic inhibitors for 30 min. The endocytic inhibitors and dosages used in this study are as follows: cytochalasin D, 2.5 μg/mL (actin/micropinocytosis inhibitor); methyl-β-cyclodextrin 2.5 mg/mL (caveolae mediated endocytsis inhibitor); nocodazole, 5 μg/mL (micropinocytosis inhibitor); Polyinosinic acid (poly I), 10 μg/mL (phagocytosis inhibitor); wortmannin, 100 ng/mL (transferrin related receptor mediated endocytosis inhibitor); dynasore, 10 μg/mL (dynamin related endocytosis). After the pre-incubation with the small molecules, 5 mol% BODIPY-lipid labeled LNPs, which encapsulated 50% Cy5-labeled mRNA and 50% normal mRNA, were added into each well of cells, at a dose of 500 ng mRNA/well. After 4 h incubation at 37 °C, the cellular uptake was determined by High Content Imaging and Analysis System (Cell Voyager CV8000, Yokogawa). Prior to imaging, the nuclei were stained with Hoechst 33342 (1 μg/mL) for 10 min. Flow cytometry was performed after High Content Imaging and intensity was applied for quantitation.

### Cytotoxicity assays

For human T cell cytotoxicity, activated T cells were transfected with 5 μg CAR muPAR-hCAR mRNA or hCD19-hCAR per million T cells using PL40 LNP, after activation. Forty hours later, the transfected T cells were used for co-culture with the target cells. NALM6 cells stably expressing GFP and NIH 3T3 cells stably expressing mouse uPAR and ZsGreen were used as target cells. Each well of a round-bottom 96-well plate contained 15,000 target cells in X-Vivo15 medium. Subsequently, T cells were added at ratios of Effector: Target (E: T) = 10: 1, using a T cell culture medium that contained 300 U/ml human recombinant IL-2.

For mouse T cell cytotoxicity, activated primary CD8^+^ T cells were transfected with 5 μg Circ muPAR-mCAR mRNA per million cells using PL40 LNPs. Twenty four hours later, the T cells were cultured with the target cells. Each well of a round-bottom 96-well plate contained 10,000 NIH 3T3-muPAR-ZsGreen cells. Subsequently, T cells were added at ratios of E: T = 40: 1, 20: 1, 10: 1.

T cells transfected Lin and Circ CAR#20 were cultured following the method mentioned above. AGS (huPAR^+^), THP-1 wild type (huPAR^+^) and THP-1 huPAR KO expressing GFP were used as target cells and non-target control cells. Each well of a round-bottom 96-well plate contained 10,000 cells in X-Vivo15 medium. Subsequently, T cells were added at ratios of E: T = 0.25:1, 1:1, and 4:1.

All the co-culturing processes were monitored by IncuCyte SX5.

### *In vivo* expression

6 to 8-week-old C57BL/6 mice were *i.v.* injected with CAMP LNPs containing mLuc (0.25 mg/kg). At 6h after injection, bioluminescence images were taken at 4 min after intraperitoneal treatment of D-Luciferin potassium salt (150 mg/kg) using an IVIS imaging system (Perkin Elmer).

### Gene editing (Cre mRNA) in the Loxp-GFP-Luciferase mice model, and flow cytometry

PL40 Cre mRNA formulation was prepared as described above. The PL40 CD3 fab formulation was prepared using the following method: The LNP ethanol phase consisted of PL40, DOPE, cholesterol, DMG-PEG2000, DSPE-PEG2000-Maleimide. The ethanol and the aqueous phase have a volume ratio of 1:3. The LNP was prepared using microfluidics, followed by overnight dialysis. CD3 fab and TCEP were reacted at 10°C for 2 hours, then mixed with the LNP and reacted overnight at 4°C. The mixture was purified using a 100kDa ultrafiltration membrane, followed by seven rounds of ultrafiltration purification with the addition of PBS in a volume ratio of 1:1. The two formulations were administrated i.v. at dose of 0.5mg/kg. After 2 d, mice were sacrificed, the liver and spleen were imaged using an IVIS Lumina system (Perkin Elmer).

Afterwards, the mouse spleen was homogenized into a single-cell suspension using a 70 μm cell strainer. ACK lysis buffer was used to remove red blood cells, followed by a single wash with PBS. The cells were then transferred to staining buffer and incubated at 4°C in the dark for 30 minutes. After one additional wash with pre-chilled PBS, the cells were resuspended in PBS for flow cytometry analysis. The following flow cytometry antibodies were used: APC anti-mouse CD3 (Biolegend), PE anti-mouse CD8a (Biolegend), PerCP Cy5.5 anti-mouse CD4 (Biolegend), BV605 anti-human/mouse CD11b (Biolegend).

### Animal models

All animal research was in compliance with ethical regulations approved by Peking University’s Institutional Animal Care and Use Committee.

### *In vivo* induction of CCl_4_-induced liver fibrosis and treatment

Liver fibrosis mouse models were established by intraperitoneal injecting 50 µl 35% CCl_4_/olive oil into 8 weeks old C57BL/6J male mice, three times per week for 5 weeks in total. Two days after final CCl_4_ injection, the first dose of LNPs (30 µg mRNA/mouse) was injected through the tail vein, and then the LNP was i.v injected every 4 days.

CD8a antibody (BioXcell, BE0061) was *i.p.* injected at 400 μg/mouse 2 days before LNPs administration.

### CIA model

An autoimmune CIA mouse model was established by the following protocol. Bovine type II collagen was dissolved in 0.05M acetic acid, then mixed with an equal volume of complete Freund’s adjuvant (CFA, 2 mg/mL), and emulsified with a homogenizer in an ice bath to get a 1mg/ml bovine type II collagen emulsion. On day 0, 7-8 weeks old DBA/1 male mice were injected subcutaneously with 100 µg of bovine type II collagen at the root of tail. On day 21, a booster immunization of 100 µg type II collagen emulsified in incomplete Freund’s adjuvant (2 mg/mL) was injected subcutaneously at the tail root.

The treatment began on day 28 after the first immunization. The CIA mice with inflammation symptoms were randomly assigned into four groups (n = 6). 30 µg muPAR-mCAR mRNA encapsulated in PL40 LNPs and the equal PL40 LNPs vehicle were administrated every 4 days via the tail vein, 5 mg/kg MTX was injected intraperitoneally every 4 days. The paw thickness was measured with caliper. The clinical scores were given by a blinded researcher based on the following criteria: 0, normal; 1, mild redness and swelling of the ankle or wrist, or distinct redness and swelling limited to digits; 2, moderate redness and swelling of ankle or wrist; 3, severe redness and swelling of the entire paw and digits; 4, Maximum redness and swelling throughout the paw and digits. The scores of each paw were added together to get the final score.

### Histological scoring of mouse arthritic joints

The scoring criteria for pathological sections are as follows. The inflammation level was assigned a score of 0-4 from weak to strong based on H&E staining, according to the criteria: 0, normal; 1, low degree of inflammatory cell infiltration in the synovial membrane area; 2, mild infiltration; 3, moderate infiltration; 4, severe infiltration. Cartilage erosion was assessed by safranin O/ Fast green staining and was assigned a score of 0-4 according to the criteria: 0, normal; 1, localized cartilage erosion; 2, more extensive cartilage erosion; 3, severe cartilage erosion; 4, erosion of the whole cartilage. The histological scoring was performed by an uninformed experimenter.

## Statistical Analysis

Data are expressed as mean ± SD as illustrated on the figure legend. Statistical significance was determined using a two-tailed unpaired Student’s t-test when only two value sets were compared or by ANOVA for comparison between multiple groups via GraphPad Prism 9.5. Exact P values are documented in the figures or figure legends. Differences were considered to be significant if P<0.05 (*P<0.05, **P<0.01, ***P<0.001, ****P<0.0001 unless otherwise indicated). For the reproducibility of the results, all the experiments were repeated independently with similar results.

## Supplementary Materials

Supplementary methods

Figs. S1 to S11

Table S1

## Acknowledgments

We thank the State Key Laboratory of Natural and Biomimetic Drugs, Peking University Biological Imaging Facility for confocal, animal, and tissue imaging services.

## Author contributions

L.M., Z.H.Z., B.M and B.Y.L. are responsible for all phases of the research. Z.H.Z, B.M., B.Y.L, Z.W.L., M.G., H.L.Z., performed experiments. J.H. and Y.Y. provided purified circular RNA. Y.W. and W.Y. helped with SAXS experiment. R.P. helped with scRNA data analysis. X.G. and R.W performed the preparation and purification of human uPAR monoclonal antibodies. Y.J.Z. helped with CD8^+^ mouse T transfection and cytotoxicity assay. Z.H.Z., B.M., B.Y.L. Y.Y.H. and L.M. wrote the manuscript. X.Q.H, B.D.C., D.M.Z., M.D. and L.M. provided conceptual advice and supervised the study. All the authors discussed the results and assisted in the preparation of the manuscript.

## Data and materials availability

All data supporting the findings of this study are presented in the main article, supplementary information, and Source Data file. The data that support the findings of this study are available from the corresponding author upon reasonable request. All the source data are provided as a “Source Data” file.

