## Supplementary Materials for "Cardiolipin-mimic lipid nanoparticles without antibody modification delivered senolytic in-vivo CAR-T therapy for inflamm-aging"

#, contributing equally,

**Supplementary Methods**

**Isolation, activation and culture of primary human T cells**

Thawed human PBMCs were used to isolate human CD3^+^ T cells which were purified using the EasySep™ Human T Cell Isolation Kit (Stemcell). T cells were cultured in X-VIVO15(Lonza) supplemented with 5% inactivated fetal bovine serum (FBS, Gibco), human recombinant IL-2 (300 U/mL, Novoprotein) and 1% penicillin/streptomycin (P/S). To activate T cells, ImmunoCult™ Human CD3/CD28 T Cell Activator (Stemcell) was added to the culture for 3 days.

**Isolation, activation and culture of primary mouse CD8^+^ T cells**

Spleens from 6-8-week-old female C57BL/6 mice were collected, mashed over a 70 μm cell strainer. Murine CD8^+^ T cells were isolated according to the instructions of EasySep Mouse T cell Isolation Kit (StemCell Techonogies,19853). T cells were plated at 10^6^/mL in 24-or 48-well plates in T cell medium (RPMI 1640-Glutmax medium supplemented with 10% FBS, 1% penicillin/streptomycin (P/S), 1 mM sodium pyruvate, 50 μM β-mercaptoethanol, 10 mM nonessential amino acids, and 50 ng/mL human IL-2) and stimulated with anti-mouse CD3/CD28 antibody-coated beads (Invitrogen) at a bead to cell ratio of 1:1. 48 h after T cell activation, the cells were transferred to each group for LNP transduction.

**Single-cell RNA-seq (scRNA-seq) analysis**

We obtained the single-cell data from the study entitled "Defining inflammatory cell states in rheumatoid arthritis joint synovial tissues by integrating single-cell transcriptomics and mass cytometry," published in 2019, and conducted subsequent analyses. Each sample underwent analysis using the Scanpy pipeline for integrated analysis. Genes expressed in fewer than 3 cells, cells expressing fewer than 200 or more than 6,000 genes (outliers), or with mitochondrial gene expression exceeding 25% were excluded. Subsequently, the harmony algorithm was applied to mitigate batch effects across samples. Principal component analysis (PCA) was performed on the top 2,000 highly variable genes. UMAP dimensional reduction was then applied to the scaled matrix using the first 50 principal components to derive a two-dimensional representation.

**Histology, immunohistochemistry (IHC) and multiplex immunohistochemistry (mIHC)**

After the mice were sacrificed, the liver of CCl_4_ induced liver fibrosis mice and the ankle joint of CIA mice were fixed in 4% paraformaldehyde and embedded in paraffin. SA-β-gal staining was performed with β-Galactosidase Staining Kit (Beyotime, China), according to the manufacturer’s instructions, and the tissue sections were counterstained with eosin. H&E staining and Safranin O/Fast green stainings performed by Servicebio. For mIHC, we followed the manufacturer’s instructions of Multitarget detection kit (Wellgene, #RD1401). Macrophages were stained with F4/80 antibody (CST, #70076, 1:500 dilution), T cells were stained with CD3ε antibody (CST, E4T1B, 1:1000 dilution), other antibodies included uPAR (Abcam, AB307895, 1:2000 dilution), α-SMA (CST, 14968S, 1:1000), CD206 (Abcam, AB300621, 1:2000).

**Animal immunization**

Six-eight-week-old Balb/C mice were immunized by injection of uPAR recombinant protein. The first booster immunization was performed 2 weeks after the initial immunization, and the second booster immunization was performed 3 weeks after the initial immunization. Blood was collected from the eye sockets one week after each booster immunization for serum immune titer detection. After diluting the obtained mouse immune serum, the specific binding of uPAR recombinant protein and uPAR overexpressing cells was detected by ELISA method, and mice with high serum titer were selected for subsequent fusion and screening.

**Fusion of specific memory B cells and hybridoma**

The spleens of mice with high serum uPAR antibody content were taken and grounded to obtain a single cell suspension. The specific memory B cells in the spleen were electrofused with mouse myeloma cells (sp2/0). The fused cells were screened using HAT medium and HT medium. The culture supernatant of the fused cells was collected for subsequent screening.

**ELISA binding assay**

In a 96-well plate, 100 μL uPAR protein (1mg/ml, diluted in PBS) was added and fixed overnight at 4℃. The protein solution was discarded, washed with PBS, and then added with 100 μL 5% skim milk to block at 37℃ for 2 hours. The plate was then washed 3 times with PBS, then added with 100 μL diluted mouse serum or monoclonal antibody (5 μg/ml) and incubated at 37℃ for 2 hours. Then, the plate was washed again with PBS, added with 100 μL anti-mouse secondary antibody, and incubated at 37℃ for 1 hour. TMB substrate was then added for color development, and finally a stop solution was added. The plate was read at OD450nm using platereader (Biotex).

**Flow cytometry analysis**

The specific binding of monoclonal culture supernatants and monoclonal antibodies to uPAR-overexpressing MDA-MB-231 cells was analyzed by flow cytometry. Briefly, 2×10^5^ cells were added to the flow tubes and blocked with 100 mg/mL IgG. The cultured supernatants or monoclonal antibodies were then added to the flow tubes and incubated on ice for 1 hour. FITC-conjugated anti-mouse secondary antibodies were then added. After washing with PBS, the fluorescence intensity of the cells was analyzed. An irrelevant mouse or IgG was used as a negative control.

**Construction and screening of uPAR truncated reporter cells**

The truncated DNA sequences of ·uPAR (D1, D2, D3, D1+D2 and D2+D3) were obtained and loaded into the pSLAP vector. Retroviruses of uPAR truncated sequences were obtained using HEK293T cells, and CT237 cells were infected with the virus to construct a stable expression cell line. Subsequently, flow cytometry was used to analyze the antigen epitopes bound by monoclonal antibodies to reporter cells.

**Affinity measurement with BLI**

For antibody affinity measurement, antibody (20 μg/mL) was loaded onto the protein G biosensors for 4 minutes. Following a short baseline in kinetics buffer, the loaded biosensors were exposed to a series of recombinant uPAR concentrations (100 nmol/L) and background subtraction was used to correct sensor drifting. All experiments were performed with shaking at 1,000 rpm. Background wavelength shifts were measured from reference biosensors that were loaded only with antibodies. ForteBio's data analysis software was used to fit the data to a 1:1 binding model to extract an association rate and dissociation rate. The Kd was calculated using the ratio k_off_/k_on_.

**Humanization of mouse monoclonal antibodies**

The CDRs in the heavy and light chains of the AB4 and AB20 antibodies were defined by a combination of three methods: Kabt, IMGT, and Paratome. The parental mouse monoclonal antibodies and the most closely related human germline sequences were then aligned. Residues known to be structurally noncritical and or not changing during in vivo maturation were identified in a mutational pedigree-guided analysis and humanized. We designed three sequences each for the heavy and light chains of the AB4 and AB20 antibodies. With different permutations, eight recombinant DNA sequences were obtained for each antibody. Finally, the sequences were loaded into the CAR construct vector and CAR-T cells expressing different permutations were prepared.

**Cardiolipin-mimic lipid synthesis**

The lipid alcohols: linoleoyl alcohol was purchased from company, and other lipid alcohols B9 and B10 synthesized using the same method. The lipids PL15, PL16, PL101, PL102, PL39, PL40, PL48, PL66 and PL68 were synthesized using the above method. The lipids PL49, PL50, PL51, PL52, PL63, PL64, PL65, PL67, PL69 and PL70 were synthesized using the same method, but different amino head groups were selected for further bromination. Structure was confirmed by ^1^H NMR,^13^C NMR spectrometry (Bruker AVANCE-400 NMR spectrometer with a Magnex Scientific superconducting magnet) and mass spectrometry (Waters Xevo G2 QTOF and ThermoFisher Orbitrap Exploris).

Synthetic route for lipid alcohol B9

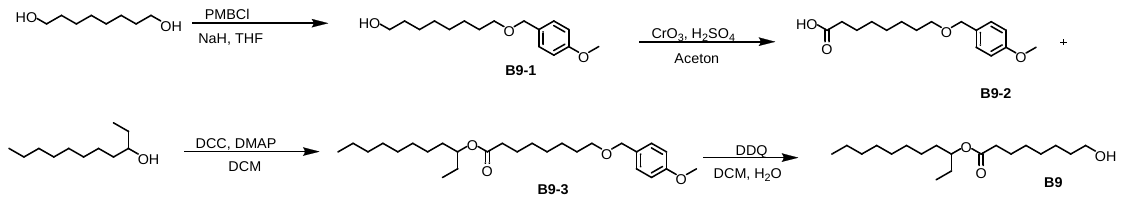

To a stirred solution of 1,8-Octanediol (2.92 g, 20 mmol) in THF (30mL) under argon was added NaH (60% dispersion in mineral oil, 1.6 g, 40 mmol). The reaction mixture was stirr for 10 min, then PMBCl (3.13 g, 20 mmol) was added. The reaction mixture was again heated to 80 °C (12 h), then allowed to cool to rt. Water was added, then the aqueous phase extracted three times with DCM. The combined organic phases were dried (Na_2_SO_4_) and concentrated. The product was purified by column chromatography, eluted with petroleum ether/ethyl acetate (25:1-5:1) to obtain the liquid compound B9-1 (3.59 g, yield 67%.) as a colourless oil. ^1^H NMR (400 MHz, CDCl_3_) δ 7.28 (d, *J*=4Hz, 2H), 6.90 (d, *J*=4Hz, 2H), 4.45(s, 2H), 3.83(s, 3H), 3.65 (t, *J*=8Hz, 2H), 3.45 (t, *J*=8Hz, 2H), 1.63-1.32(m, 12H).

To a stirred solution of alcohol B9-1 (2.66 g, 10 mmol) in acetone (30 mL) at 0 °C was added Jones' reagent (15 mL). The reaction mixture was stirred at rt for 2 h, then the reaction was quenched by addition of propan-2-ol (15 mL), and then carefully neutralised with NaHCO_3_. The solution was then extracted three times with DCM (3 x 200 mL). The combined organic layers were dried (Na_2_SO_4_) and concentrated. The product was purified by column chromatography, eluted with petroleum ether/ethyl acetate (25:1-5:1) to obtain acid B9-2 (2.44 g, yield 87%) as a white solid. ^1^H NMR (400 MHz, CDCl_3_) δ 7.28 (d, *J*=4Hz, 2H), 6.90 (d, *J*=4Hz, 2H), 4.45(s, 2H), 3.83(s, 3H), 3.45 (t, *J*=8Hz, 2H), 3.36 (t, *J*=8Hz, 2H), 1.67-1.60(m, 4H). 1.40-1.35(m, 6H).

To a stirred solution of undecan-3-ol (1.38 g, 8 mmol) and acid B9-2 (2.24 g, 8 mmol) in DCM (35 mL) was added DCC (2.47 g, 12 mmol) and DMAP (147 mg, 1.2 mmol). The reaction mixture was stirred at rt for 16 h, then vacuum suction filtration, filtrate was added with water. The layers were separated, and the aqueous layer extracted three times with DCM. The combined organic phases were washed with saturated citric acid aqueous solution, then dried (Na_2_SO_4_) and concentrated. The product was sample purified by column chromatography, elute with petroleum ether/ethyl acetate (25:1) to obtain crude ester B9-3 as a colorless oil. The crude alcohol was used in the next step without further purification.

To a stirred solution of PMB ether B9-3 in DCM (15 mL) and water (1.5 mL) was added DDQ (1.816 g, 8 mmol). The reaction mixture was stirred for 2 h, then it was quenched by addition of saturated NaHCO_3_. The layers were separated, and the aqueous phase extracted three times with DCM. The combined organic phases were washed with water three times, then dried (Na_2_SO_4_) and concentrated. The residue was purified by column chromatography (25:1 petroleum ether/ethyl acetate) to afford lipid alcohol B9 (1.63g, 64%) as a colorless oil. ^1^H NMR (400 MHz, CDCl_3_) δ 4.86-4.80 (m, 1H), 3.98 (t, *J*=8Hz, 2H), 2.31 (t, *J*=8Hz, 2H), 1.69-1.51(m, 8H), 1.36-1.28 (m, 18H), 0.91-0.87(m,6H).

Synthetic route for lipid alcohol B10

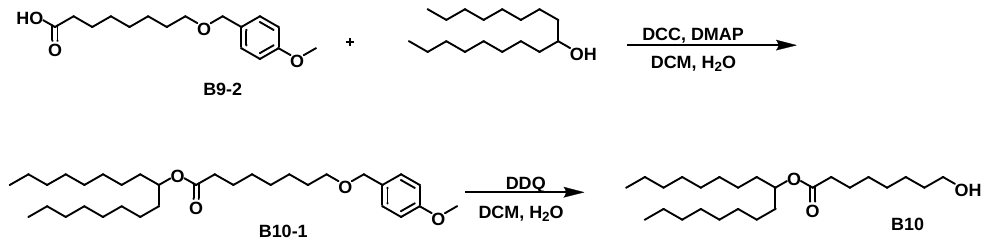

To a stirred solution of heptadecan-9-ol (2.05 g, 8 mmol) and acid B9-2 (2.24 g, 8 mmol) in DCM (35 mL) was added DCC (2.47 g, 12 mmol) and DMAP (147 mg, 1.2 mmol). The reaction mixture was stirred at rt for 16 h, then vacuum suction filtration, filtrate was added with water. The layers were separated, and the aqueous layer extracted three times with DCM. The combined organic phases were washed with saturated citric acid aqueous solution, then dried (Na_2_SO_4_) and concentrated. The product was sample purified by column chromatography, eluted with petroleum ether/ethyl acetate (25:1) to obtain crude ester B10-1 as a colorless oil. The crude alcohol was used in the next step without further purification.

To a stirred solution of PMB ether B10-1 in DCM (15 mL) and water (1.5 mL) was added DDQ (1.816 g, 8 mmol). The reaction mixture was stirred for 2 h, then it was quenched by addition of saturated NaHCO_3_. The layers were separated, and the aqueous phase extracted three times with DCM. The combined organic phases were washed with water three times, then dried (Na_2_SO_4_) and concentrated. The residue was purified by column chromatography, eluted with petroleum ether/ethyl acetate (25:1-15:1) to obtain lipid alcohol B10 (2.36g, 74%) as a colorless oil. ^1^H NMR (400 MHz, CDCl_3_): δ 4.86-4.80 (m, 1H), 3.98 (t, *J*=8Hz, 2H), 2.31 (t, *J*=8Hz, 2H), 1.69-1.51(m, 8H), 1.36-1.28 (m, 30H), 0.91-0.87(m,6H).

Synthetic route for lipids PL15, PL16, PL101, PL102, PL39, PL40, PL48, PL66 and PL68

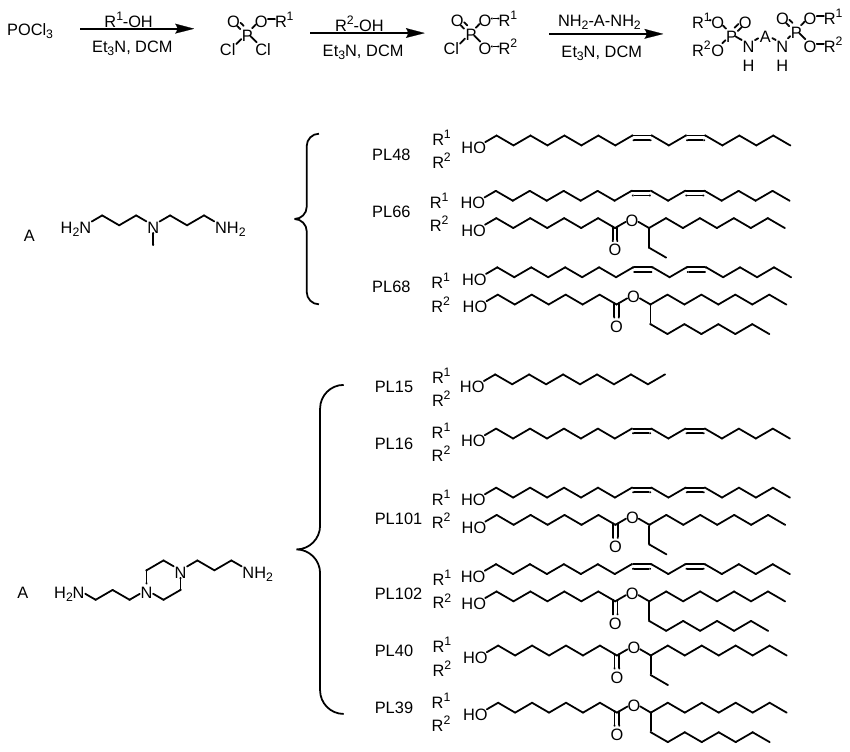

To a stirred solution of POCl_3_ (156 mg, 1 mmol) in DCM (10 mL) was added a DCM solution of R^1^-OH (1 mmol) with Et_3_N (101 mg, 1 mmol). The reaction mixture was stirred at 4 ℃ for 3 h, then the DCM solution of R^2^-OH (1 mmol) and Et_3_N (101 mg, 1 mmol) was added slowly dropwise to system and stirred at rt for 2h. Subsequently, the DCM solution of NH_2_-A-NH_2_ (1 mmol) and Et_3_N (202 mg, 2 mmol) was added slowly dropwise to system, and then stirred at rt for 2h. After completion of the reaction, wash with saturated Sodium chloride, and concentrate under reduced pressure to obtain the crude product. Purified by column chromatography, eluted with dichloromethane: methanol (30:1-15:1) to obtain the liquid compound.

Compound **PL15**: yield 87%. ^1^H NMR (400 MHz, CDCl_3_) δ 3.91-3.99 (m, 8H), 3.64-3.61 (m, 2H), 3.01-2.94(m, 4H), 2.88-2.23 (m, 12H), 1.68-1.61 (m, 12H), 1.37-1.25 (m, 64H), 0.87 (t, *J*=8Hz, 12H). ^13^C NMR (100 MHz, CDCl_3_) δ 66.41, 56.76, 53.14, 40.88, 31.92, 30.48, 30.41, 29.62, 29.58, 29.35, 29.25, 25.63, 22.69, 14.12.

Compound **PL16**: yield 80%. ^1^H NMR (400 MHz, CDCl_3_) δ 5.41-5.29 (m, 16H), 4.02-3.90 (m, 8H), 3.64 (s, 2H), 2.99-2.97(m, 4H), 2.78 (t, *J*=4Hz, 8H), 2.61-2.43 (m, 12H), 2.07-2.02 (m ,16H), 1.69-1.62 (m, 12H), 1.39-1.26 (m, 64H), 0.89 (t, *J*=8Hz, 12H). ^13^C NMR (100 MHz, CDCl_3_) δ 130.43, 130.27, 128.26, 128.13, 66.29, 56.8, 53.1, 40.85, 31.51, 30.46, 30.39, 29.65, 29.46, 29.33, 29.24, 29.21, 27.21, 27.18, 25.61, 22.56, 14.06.

Compound **PL101**: yield 62%. ^1^H NMR (400 MHz, CDCl_3_) δ 5.44-5.31 (m, 8H), 4.86-4.80 (m, 2H), 4.02-3.94 (m, 8H), 3.00-2.29 (m, 20H), 2.10-2.04(m ,8H), 1.71-1.50 (m, 24H), 1.38-1.28 (m, 68H), 0.93-0.87 (m, 18H). ^13^C NMR (100 MHz, CDCl_3_) δ 173.89, 130.50, 130.34, 128.31, 128.18, 75.55, 66.59, 56.92, 53.62, 40.85, 34.91, 33.91, 32.14, 31.80, 30.76, 30.69, 29.94, 29.81, 29.78, 29.76, 29.62, 29.54, 29.51, 29.36, 29.19, 27.50, 27.25, 25.91, 25.74, 25.61, 25.35,22.94, 22.85, 14.38, 14.35, 9.89. MS (ESI, m/z) [M+H] ^+^ calcd. For C_84_H_162_N_4_O_10_P_2_, 1149.17, found:1450.17.

Compound **PL102**: yield 56%. ^1^H NMR (400 MHz, CDCl_3_) δ 5.44-5.32 (m, 8H), 4.90-4.85 (m, 2H), 4.04-3.86 (m, 8H), 3.00-2.28 (m, 20H), 2.10-2.05(m ,8H), 1.75-1.50 (m, 24H), 1.40-1.28 (m, 92H), 0.93-0.88 (m, 18H). ^13^C NMR (100 MHz, CDCl_3_) δ 173.82, 130.47, 130.40, 128.29, 128.16, 74.41, 66.62, 56.88, 53.01, 40.78, 34.90, 34.39, 32.11, 32.78, 30.73, 30.67, 29.92, 29.78, 29.75, 29.60, 29.52, 29.49, 29.36, 29.19, 27.48, 27.46, 25.88, 25.73, 25.57, 25.32, 22.92, 22.83, 14.36, 14.33. MS (ESI, *m/z*) [M+2H] ^+^ calcd. For C_96_H_186_N_4_O_10_P_2_, 1617.36, found:809.68.

Compound **PL39**: yield 59%. ^1^H NMR (400 MHz, CDCl_3_) δ 4.89-4.86 (m, 4H), 4.01-3.93 (m, 8H), 3.05-2.27(m, 24H), 1.67-1.50 (m ,36H), 1.36-1.27 (m, 120H), 0.89 (t, *J*=8Hz, 24H). ^13^C NMR (100 MHz, CDCl_3_) δ 173.85, 74.44, 66.80, 56.82, 53.69, 40.28, 34.91, 34.39, 32.13, 30.68, 30.61, 29.81, 29.77, 29.51, 29.37, 29.21, 25.72, 25.59, 25.33, 22.94, 14.38.

Compound **PL40**: yield 46%. ^1^H NMR (400 MHz, CDCl_3_) δ 4.86-4.79 (m, 4H), 4.00-3.94 (m, 8H), 3.15-2.10(m, 24H), 1.69-1.51 (m, 36H), 1.36-1.28 (m, 72H), 0.91-0.87 (m, 24H). ^13^C NMR (100 MHz, CDCl_3_) δ 173.89, 75.54, 66.92, 56.94, 53.68, 40.81, 34.86, 33.87, 32.10, 30.59, 30.52, 29.78, 29.74, 29.48, 29.31, 29.13, 27.21, 25.66, 25.58, 25.30, 22.91, 14.35, 9.86. MS (ESI, *m/z*) [M+2H] ^+^ calcd. For C_86_H_170_N_4_O_14_P_2_, 1545.21, found:773.61.

Compound **PL48**: yield 49%. ^1^H NMR (400 MHz, CDCl_3_) δ 5.44-5.28 (m, 16H), 4.01-3.94 (m, 8H), 3.12-2.63 (m, 16H), 2.09-2.04 (m ,19H), 1.69-1.64 (m, 12H), 1.39-1.27 (m, 64H), 0.92 (t, *J*=8Hz, 12H). **^1^**^3^C NMR (100 MHz, CDCl_3_) δ 130.43, 130.27, 128.26, 128.13, 66.59, 53.66, 45.42, 40.17, 31.75, 30.71, 30.64, 29.90, 29.74, 29.57, 29.51, 29.47, 27.46, 27.42, 25.85, 22.80, 14.30

Compound **PL66**: yield 63%. ^1^H NMR (400 MHz, CDCl_3_) δ 5.44-5.31 (m, 8H), 4.88-4.79 (m, 2H), 4.02-3.83 (m, 8H), 3.24-2.75 (m, 12H), 2.32-2.27 (m, 4H), 2.09-2.02(m ,11H), 1.70-1.50 (m, 24H), 1.38-1.28 (m, 68H), 0.93-0.87 (m, 18H). ^13^C NMR (100 MHz, CDCl_3_) δ 173.85, 130.45, 130.28, 128.28, 128.14, 75.52, 66.83, 53.67, 45.93, 40.94, 34.87, 33.87, 32.10, 31.77, 31.48, 30.69, 30.62, 30.56, 29.90, 29.77, 29.74, 29.71, 29.59, 29.50, 29.48, 29.46, 29.31, 29.14, 27.47, 27.44, 27.22, 26.23, 25.87, 25.85, 25.69, 25.57, 25.30, 22.90, 22.81,14.35, 14.32, 9.86.

Compound **PL68**: yield 62%. ^1^H NMR (400 MHz, CDCl_3_) δ 5.43-5.31 (m, 8H), 4.09-3.96 (m, 12H), 3.28-2.77 (m, 12H), 2.34-2.30 (m, 2H), 2.09-2.04(m ,11H), 1.70-1.57 (m, 16H), 1.43-1.24 (m, 92H), 0.92-0.89 (m, 18H). ^13^C NMR (100 MHz, CDCl_3_) δ 176.96, 130.50, 130.29, 128.32, 128.16, 74.47, 66.93, 53.88, 45.96, 38.75, 32.76, 32.71, 32.13, 31.96, 31.79, 29.95, 29.84, 29.79, 29.72, 29.62, 29.57, 29.53, 29.49, 29.02, 28.90, 27.72, 27.68, 27.53,

27.46, 26.07, 25.90, 25.56, 22.93, 22.86, 22.84, 14.38, 14.34.

Synthetic route for lipids PL49, PL50, PL51, PL52, PL63, PL64, PL65, PL67, PL69 and PL70

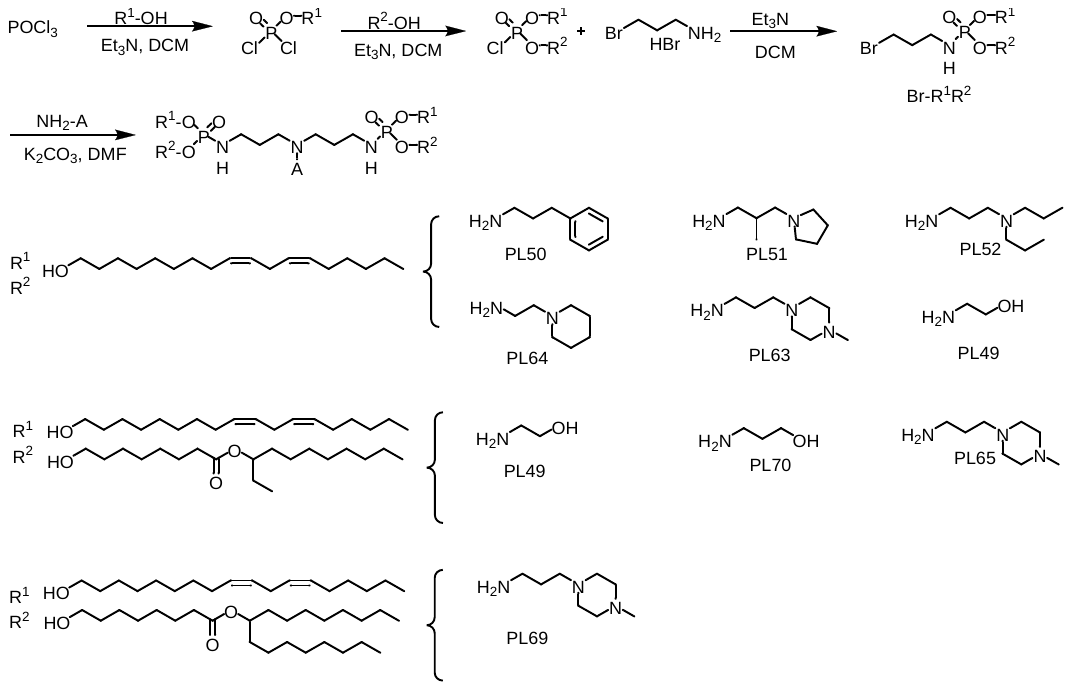

To a stirred solution of POCl_3_ (156 mg, 1 mmol) in DCM (10 mL) was added a DCM solution of R^1^-OH (1 mmol) with Et_3_N (101 mg, 1 mmol). The reaction mixture was stirred at 4℃ for 3 h, then the DCM solution of R^2^-OH (1 mmol) and Et_3_N (101 mg, 1 mmol) was added slowly dropwise to system and stirred at rt for 2h. Subsequently, the DCM solution of NH_2_-A (2 mmol) and Et_3_N (202 mg, 2 mmol) was added slowly dropwise to system, and then stirred at rt for 2h. After completion of the reaction, wash with saturated Sodium chloride, and concentrate under reduced pressure to obtain the crude product. Purify by column chromatography, elute with dichloromethane: methanol (60:1-30:1) to obtain the liquid compound **Br-R^1^R^2^**.

To a solution of Br-R^1^R^2^ (1 mmol) in DMF (10 mL) was added A-NH_2_ (0.5 mmol) and K_2_CO_3_ (138 mg,1 mmol), then the reaction mixture was stirred at rt for 24 h. After completion of the reaction, wash with saturated Sodium chloride, and concentrate under reduced pressure to obtain the crude product. Purify by column chromatography, elute with dichloromethane: methanol 25:1-15:1 to obtain the liquid compound.

Compound **PL49**: yield 58%. ^1^H NMR (400 MHz, CDCl_3_) δ 5.42-5.32 (m, 16H), 4.01-3.97 (m, 8H), 3.76-3.67 (m, 2H), 3.51-3.41 (m, 2H), 3.04-2.91 (m, 4H), 2.79 (t, *J*=4Hz, 8H), 2.71-2.67 (m, 6H), 2.10-2.05 (m ,16H), 1.92-1.81 (m, 4H), 1.72-1.65 (m, 8H), 1.40-1.28 (m, 64H), 0.91 (t, *J*=4Hz, 12H). ^13^C NMR (100 MHz, CDCl_3_) δ 130.44, 130.28, 128.28, 128.13, 66.72, 58.71, 56.24, 52.03, 39.88, 31.75, 30.70, 30.64, 29.90, 29.73, 29.58, 29.51, 29.47, 27.46, 27.43, 25.86, 22.80, 14.30

Compound **PL50**: yield 63%. ^1^H NMR (400 MHz, CDCl_3_) δ 7.31-7.27 (m, 2H), 7.21-7.18 (m, 3H), 5.42-5.31 (m, 16H), 4.01-3.92 (m, 8H), 3.13-2.45 (m, 20H), 2.09-2.04 (m ,16H), 1.92-1.80 (m, 2H), 1.68-1.63 (m, 12H), 1.39-1.28 (m, 64H), 0.91 (t, *J*=4Hz, 12H). ^13^C NMR (100 MHz, CDCl_3_) δ 130.39, 130.21, 128.82, 128.74, 128.52, 128.22, 128.08, 66.84, 56.24, 53.46, 40.07, 32.90, 31.70, 31.64, 30.64, 30.57, 29.86, 29.69, 29.52, 29.47, 29.43, 27.41, 27.37, 25.81, 22.75, 14.26.

Compound **PL51**: yield 67%. ^1^H NMR (400 MHz, CDCl_3_) δ 5.42-5.31 (m, 16H), 4.04-3.92 (m, 8H), 3.08-2.94 (m, 10H), 2.84-2.63 (m, 12H), 2.09-2.04 (m ,18H), 1.92-1.57 (m, 16H), 1.39-1.28 (m, 65H), 1.02 (t, *J*=8Hz, 3H), 0.91 (t, *J*=4Hz, 12H). ^13^C NMR (100 MHz, CDCl_3_) δ 130.43, 130.27, 128.17, 128.12, 66.66, 61.43, 60.23, 56.69, 52.57, 40.05, 31.76, 31.69, 30.68, 30.61, 29.89, 29.70, 29.58, 29.49, 29.45, 27.46, 27.43, 26.21, 25.87, 23.37, 22.81, 15.87, 14.31.

Compound **PL52**: yield 71%. ^1^H NMR (400 MHz, CDCl_3_) δ 5.42-5.30 (m, 16H), 4.02-3.94 (m, 8H), 3.00-2.93 (m, 4H), 2.80-2.46 (m, 20H), 2.08-2.03 (m, 16H), 1.82-1.59 (m, 18H), 1.39-1.20 (m, 64H), 0.96-0.88 (m, 18H)。^13^C NMR (100 MHz, CDCl3) δ 130.46, 130.29, 128.28, 128.15, 66.59, 56.45, 52.29, 40.59, 31.77, 30.76, 30.69, 29.93, 29.76, 29.60, 29.53, 29.51, 27.48, 27.45, 25.89, 22.82, 14.32, 11.91.

Compound **PL63**: yield 76%. ^1^H NMR (400 MHz, CDCl_3_) δ 5.41-5.31 (m, 16H), 4.02-3.93 (m, 8H), 3.28-3.24 (m, 4H), 3.05-2.91 (m, 6H), 2.80-2.54 (m, 21H), 2.09-2.04 (m, 16H), 1.82-1.64 (m, 14H), 1.39-1.27 (m, 64H), 0.91 (t, *J*=4Hz, 12H). ^13^C NMR (100 MHz, CDCl_3_) δ 130.48, 130.30, 128.27, 128.15, 66.67, 62.43, 56.51, 54.83, 52.69, 45.75, 40.25, 31.76, 30.68, 30.61, 29.89, 29.70, 29.58, 29.49, 29.45, 27.46, 27.43, 26.21, 25.87, 22.81, 14.31.

Compound **PL64**: yield 73%. ^1^H NMR (400 MHz, CDCl_3_) δ 5.44-5.31 (m, 16H), 4.02-3.94 (m, 8H), 3.04-2.97 (m, 4H), 2.79 (t, *J*=4Hz, 8H), 2.52-2.46 (m, 12H), 2.09-2.04 (m ,16H), 1.72-1.64 (m, 16H), 1.41-1.27 (m, 66H), 0.91 (t, *J*=4Hz, 12H). ^13^C NMR (100 MHz, CDCl_3_) δ 130.48, 130.33, 128.28, 128.16, 66.71, 56.55, 54.54, 51.31, 40.39, 31.77, 30.64, 29.92, 29.75, 29.59, 29.52, 27.48, 27.45, 25.88, 25.86, 22.82, 14.33.

Compound **PL65**: yield 48%. ^1^H NMR (400 MHz, CDCl_3_) δ 5.41-5.32 (m, 8H), 4.86-4.29 (m, 2H), 4.02-3.93 (m, 8H), 3.38-3.25 (m, 4H), 3.03-2.97 (m, 4H), 2.79 (t, *J*=4Hz, 4H), 2.59-2.49(m ,12H), 2.38(s ,3H), 2.30 (t, *J*=4Hz, 4H), 2.09-2.04(m ,8H), 1.80-1.52 (m, 26H), 1.36-1.27 (m, 68H), 0.92-0.87 (m, 18H). ^13^C NMR (100 MHz, CDCl_3_) δ 173.87, 130.46, 130.30, 128.27, 128.15, 75.51, 66.60, 62.20, 56.76, 55.05, 52.93, 45.93, 40.94, 34.87, 33.87, 32.10, 31.77, 31.48, 30.69, 30.62, 30.56, 29.90, 29.77, 29.74, 29.71, 29.59, 29.50, 29.48, 29.46, 29.31, 29.14, 27.47, 27.44, 27.22, 26.23, 25.87, 25.85, 25.69, 25.57, 25.30, 22.90, 22.81,14.35, 14.32, 9.86.

Compound **PL67**: yield 69%. ^1^H NMR (400 MHz, CDCl_3_) δ 5.42-5.31 (m, 8H), 4.84-4.80 (m, 2H), 4.01-3.96 (m, 8H), 3.86-3.50 (m, 4H), 3.03-2.72 (m, 14H), 2.10-2.05(m, 8H), 1.72-1.51 (m, 24H), 1.38-1.28 (m, 68H), 0.93-0.87 (m, 18H). ^13^C NMR (100 MHz, CDCl_3_) δ 173.91, 130.51, 130.53, 128.32, 128.19, 75.57, 66.87, 58.74, 56.25, 52.06, 39.98, 34.90, 33.91, 32.14, 31.81, 30.75, 30.69, 30.61, 29.97, 29.82, 29.79, 29.63, 29.59, 29.56, 29.52, 29.38, 29.22, 27.52, 27.48, 27.24, 25.92, 25.76, 25.62, 25.35, 22.95, 22.86,14.39, 14.36, 9.89.

Compound **PL69**: yield 73%. ^1^H NMR (400 MHz, CDCl_3_) δ 5.44-5.31 (m, 8H), 4.08 (t, *J*=8Hz, 4H), 4.01-3.95 (m, 8H), 3.30-3.25 (m, 4H), 3.04-2.90 (m, 6H), 2.79 (t, *J*=8Hz, 4H), 2.62-2.52 (m, 10H), 2.40 (s, 3H), 2.35-2.30(m ,2H), 2.09-2.04(m ,8H), 1.82-1.55 (m, 26H), 1.42-1.27 (m, 84H), 0.93-0.88 (m, 18H). ^13^C NMR (100 MHz, CDCl_3_) δ 176.93, 130.46, 130.30, 128.28, 128.15, 75.52, 66.62, 62.20, 56.71, 54.99, 52.86, 46.06, 40.44, 32.76, 32.10, 31.94, 31.77, 31.48, 30.70, 30.63, 30.53, 29.90, 29.80, 29.71, 29.69, 29.59, 29.50, 29.46, 28.87, 27.71, 27.66, 27.47, 27.44, 26.22, 25.87, 25.85, 25.54, 22.90, 22.83, 22.82, 14.35, 14.32.

Compound **PL70**: yield 57%. ^1^H NMR (400 MHz, CDCl_3_) δ 5.44-5.29 (m, 8H), 4.86-4.79 (m, 2H), 4.01-3.94 (m, 8H), 3.78-3.76 (m, 2H), 3.49-3.45 (m, 2H), 3.01-2.77 (m, 14H), 2.30 (t, *J*=8Hz, 4H), 2.09-2.04(m ,8H), 1.87-1.51 (m, 26H), 1.37-1.28 (m, 68H), 0.93-0.87 (m, 18H). ^13^C NMR (100 MHz, CDCl_3_) δ 173.91, 130.49, 130.33, 128.30, 128.18, 75.56, 66.77, 56.96, 53.69, 51.57, 39.65, 34.89, 33.89, 32.13, 31.79, 30.74, 30.67, 30.59, 29.94, 29.80, 29.77, 29.61, 29.55, 29.51, 29.34, 29.18, 27.50, 27.47, 27.23, 25.90, 25.72, 25.60, 25.33, 22.93, 22.84,14.37, 14.34, 9.88.

Synthetic route for lipid PL62

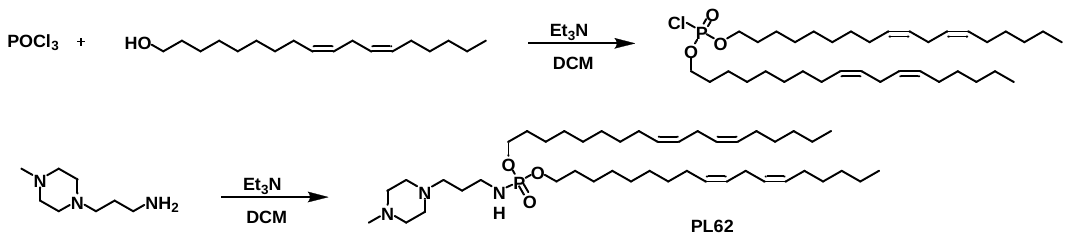

To a stirred solution of POCl_3_ (156 mg, 1 mmol) in DCM (10 mL) was added a DCM solution of (9Z,12Z)-octadeca-9,12-dien-1-ol (532 mg, 2 mmol) with Et_3_N (202 mg, 2 mmol). The reaction mixture was stirred at rt for 3 h, then the DCM solution of 3-(4-methylpiperazin-1-yl)propan-1-amine (157 mg, 1 mmol) and Et_3_N (101 mg, 1 mmol) was added slowly dropwise to system, and then stirred at rt for 2h. After completion of the reaction, wash with saturated Sodium chloride, and concentrate under reduced pressure to obtain the crude product. Purify by column chromatography, elute with dichloromethane: methanol (30:1-15:1) to obtain the liquid compound PL62 (614 mg, 83.6%).^1^H NMR (400 MHz, CDCl_3_) δ 5.42-5.29 (m, 8H), 4.01-3.95 (m, 4H), 3.04-2.49 (m, 16H), 2.36 (S, 3H), 2.10-2.04 (m, 8H), 1.72-1.65 (m, 6H), 1.42-1.26 (m, 32H), 0.90 (t, *J*=8Hz 6H). ^13^C NMR (100 MHz, CDCl_3_) δ 130.50, 130.33, 128.31, 128.18, 66.59, 57.10, 55.13, 53.10, 45.97, 41.15, 31.80, 30.76, 30.69, 29.93, 29.75, 29.62, 29.53, 29.50, 27.50, 27.47, 25.91, 22.85, 14.35.

Synthetic route for lipid PL71

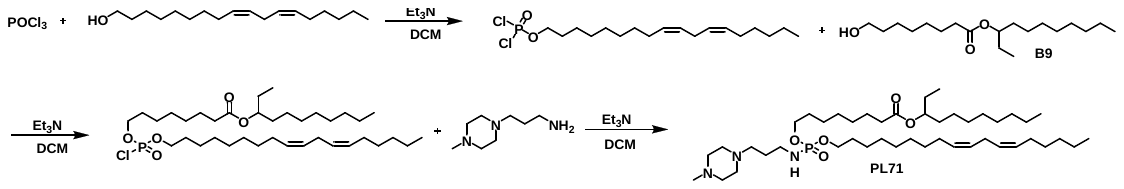

To a stirred solution of POCl_3_ (156 mg, 1 mmol) in DCM (10 mL) was added a DCM solution of (9Z,12Z)-octadeca-9,12-dien-1-ol (266 mg, 1 mmol) with Et_3_N (101 mg, 1 mmol). The reaction mixture was stirred at 4℃ for 3 h, then the DCM solution of B9 (314 mg, 1 mmol) and Et_3_N (101 mg, 1 mmol) was added slowly dropwise to system and stirred at rt for 2h. Subsequently, the DCM solution of 3-(4-methylpiperazin-1-yl) propan-1-amine (157 mg, 1 mmol) and Et_3_N (101 mg, 1 mmol) was added slowly dropwise to system, and then stirred at rt for 2h. After completion of the reaction, wash with saturated Sodium chloride, and concentrate under reduced pressure to obtain the crude product. Purify by column chromatography, elute with dichloromethane: methanol (30:1-15:1) to obtain the liquid compound PL71 (682.17 mg, 87.21%).^1^H NMR (400 MHz, CDCl_3_) δ 5.43-5.32 (m, 4H), 4.84-4.81 (m, 1H), 4.01-3.94 (m, 4H), 3.03-2.99 (m, 2H), 2.79 (t, *J*=8Hz, 2H), 2.52-2.39 (m, 10H), 2.32-2.28 (m, 5H), 2.09-2.04 (m, 4H), 1.71-1.50 (m, 12H), 1.38-1.27 (m, 34H), 0.92-0.87 (m, 9H). ^13^C NMR (100 MHz, CDCl_3_) δ 173.88, 130.47, 130.31, 128.28, 128.16, 75.53, 66.51, 57.14, 55.24, 53.26, 46.08, 41.18, 34.88, 33.88, 32.10, 31.77, 30.73, 30.66, 29.91, 29.78, 29.75, 29.72, 29.59, 29.51, 29.48, 29.33, 29.16, 27.73, 27.67, 27.47, 27.45, 27.22, 25.88, 25.72, 25.58, 25.32, 22.91, 22.82, 14.35, 14.32, 9.86.

Synthetic route for lipid PL82

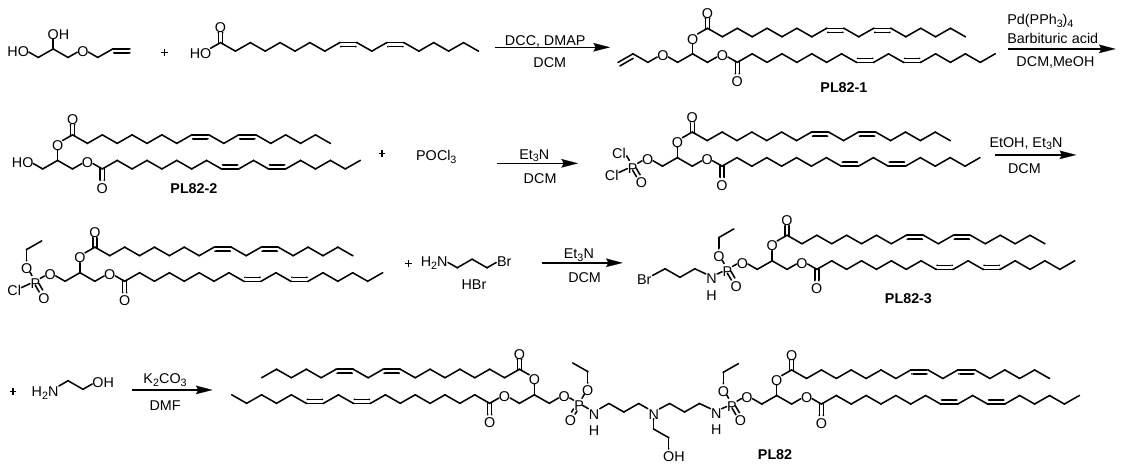

To a stirred solution of 3-allyloxy-1,2-propanediol (660 mg, 5 mmol) in DCM (10 mL) was added a DCM solution of linoleic acid (700 mg, 2.5 mmol) with DCC (515 mg, 2.5 mmol) and DMAP (30mg, 0.25 mmol), then stirred at 4℃ for 16 h. After completion of the reaction, wash with saturated Sodium chloride, and concentrate under reduced pressure to obtain the crude product. Purify by column chromatography, elute with petroleum ether: ethyl acetate (25:1) to obtain the liquid compound **PL82-1** (2.95 g, 90% )。^1^H NMR (400 MHz, CDCl_3_): 5.94-5.84 (m, 1H), 5.44-5.19 (m, 11H), 4.38-4.34 (m, 1H), 4.22-4.17 (m, 1H), 4.03-4.00 (m, 2H), 3.58 (d, *J*=4Hz, 2H), 2.79 (t, *J*=4Hz, 4H),2.37-2.30 (m, 4H), 2.10-2.04 (m, 8H), 1.67-1.60 (m, 4H),1.41-1.27 (m, 28H), 0.89 ( t, *J*=8Hz, 6H).

To a stirred solution of PL82-1 (2.95 g, 4.5 mmol) in DCM/MeOH (5 ml/5 mL) was added Pd(PPh3)4 (1155 mg, 1 mmol) and barbituric acid (768 mg, 6 mmol), then stirred at 60℃ for 16 h. After completion of the reaction, wash with saturated Sodium chloride, and concentrate under reduced pressure to obtain the crude product. Purify by column chromatography, elute with petroleum ether: ethyl acetate (10:1) to obtain the liquid compound **PL82-2**(2.21 g, 79.7%)。^1^H NMR (400 MHz, CDCl_3_) δ 5.44-5.31 (m, 8H), 5.13-5.08 (m, 1H), 4.36-4.13 (m, 2H), 3.76-3.74 (m, 2H), 2.79 (t, *J*=4Hz, 4H), 2.39-2.33 (m, 4H), 2.10-2.05 (m, 8H), 1.67-1.62 (m, 4H), 1.40-1.27(m, 28H), 0.92 (t, *J*=8Hz, 6H).

To a stirred solution of POCl_3_ (156 mg, 1 mmol) in DCM (10 mL) was added a DCM solution of PL82-2 (616 mg, 1 mmoL) with Et_3_N (101 mg, 1 mmol). The reaction mixture was stirred at 4℃ for 3 h, then the DCM solution of ethanol (46 mg, 1 mmol) and Et_3_N (101 mg, 1 mmol) was added slowly dropwise to system and stirred at rt for 2h. Subsequently, the DCM solution of 3-bromopropylamine hydrobromide (218 mg, 1 mmol) and Et_3_N (101 mg, 1 mmol) was added slowly dropwise to system, and then stirred at rt for 2h. After completion of the reaction, wash with saturated Sodium chloride, and concentrate under reduced pressure to obtain the crude product. Purify by column chromatography, elute with dichloromethane: methanol (30:1) to obtain the liquid compound **PL 82-3** (657.54 mg, 68%)。^1^H NMR (400 MHz, CDCl_3_) δ 5.44-5.45 (m, 9H), 4.38-4.06 (m, 6H), 3.51-3.48 (m, 2H), 3.15-3.07 (m, 2H), 2.79 (t, *J*=4Hz, 4H), 2.38-2.32 (m, 4H), 2.10-2.03(m, 10H), 1.65-1.61 (m, 4H), 1.40-1.27 (m, 31H), 0.91 (t, *J*=4Hz, 6H).

To a solution of PL82-3 (843.48 mg，1 mmol) in DMF (10 mL) was added ethanolamine (30.5 mg，0.5 mmol) and K_2_CO_3_ (138 mg,1 mmol), then the reaction mixture was stirred at rt for 24 h. After completion of the reaction, wash with saturated Sodium chloride, and concentrate under reduced pressure to obtain the crude product. Purify by column chromatography, elute with dichloromethane: methanol (30:1) to obtain the liquid compound **PL82** (373.18mg, 47%)。^1^H NMR (400 MHz, CDCl_3_) δ 5.44-5.24 (m, 18H), 4.38-4.04 (m, 12H), 3.81-3.66 (m, 2H), 3.21-2.75 (m, 18H), 2.37-2.31 (m, 8H), 2.09-2.04(m,16H), 1.92-1.85 (m, 4H),1.65-1.59 (m, 8H), 1.40-1.28 (m, 62H), 0.91 (t, *J*=4Hz, 12H). ^13^C NMR (100 MHz, CDCl_3_) δ 173.62, 173.23, 130.48, 130.24, 130.21, 128.34, 128.32, 128.13, 69.99, 64.11, 63.05, 62.31, 56.02, 52.09, 39.96, 34.45, 34.28, 32.15, 31.76, 29.88, 29.77, 29.59, 29.48, 29.46, 29.39, 29.35, 29.33, 27.44, 25.87, 25.11, 25.08, 22.82, 16.50, 14.32.

Synthetic route for lipid PL86

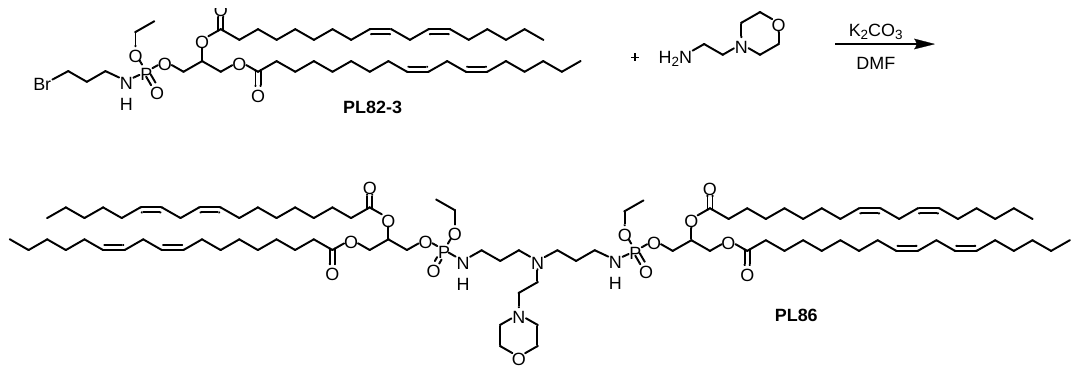

To a solution of PL82-3 (843.48 mg，1 mmol) in DMF (10 mL) was added 4-(2-aminoethyl)morpholine (65 mg，0.5 mmol) and K_2_CO_3_ (138 mg,1 mmol), then the reaction mixture was stirred at rt for 24 h. After completion of the reaction, wash with saturated Sodium chloride, and concentrate under reduced pressure to obtain the crude product. Purify by column chromatography, elute with dichloromethane: methanol (30:1) to obtain the liquid compound **PL86** (613.46mg, 74%)。^1^H NMR (400 MHz, MeOD) δ 5.44-5.24 (m, 18H), 4.36-4.04 (m,12H), 3.84-3.66 (m, 4H), 3.40-3.24 (m, 4H), 3.05-2.99 (m, 4H), 2.79 (t, *J*=4Hz, 8H), 2.62-2.41 (m, 8H), 2.37-2.31 (m, 8H), 2.09-2.04(m, 16H), 1.83-1.80 (m, 4H), 1.65-1.60 (m, 8H),1.41-1.26 (m, 62H), 0.93-0.87 (m, 12H). ^13^C NMR (100 MHz, CDCl_3_) δ 173.56, 173.11, 130.48, 130.24, 130.22, 128.34, 128.32, 128.14, 69.86, 67.06, 64.26, 62.95, 62.27, 57.82, 53.68, 53.63, 38.58, 34.44, 34.27, 31.77, 29.87, 29.77, 29.59, 29.46, 29.45, 29.38, 29.35, 29.32, 27.44, 25.87, 25.11, 25.08, 22.82, 16.49, 14.32.

**Supplementary Figures**

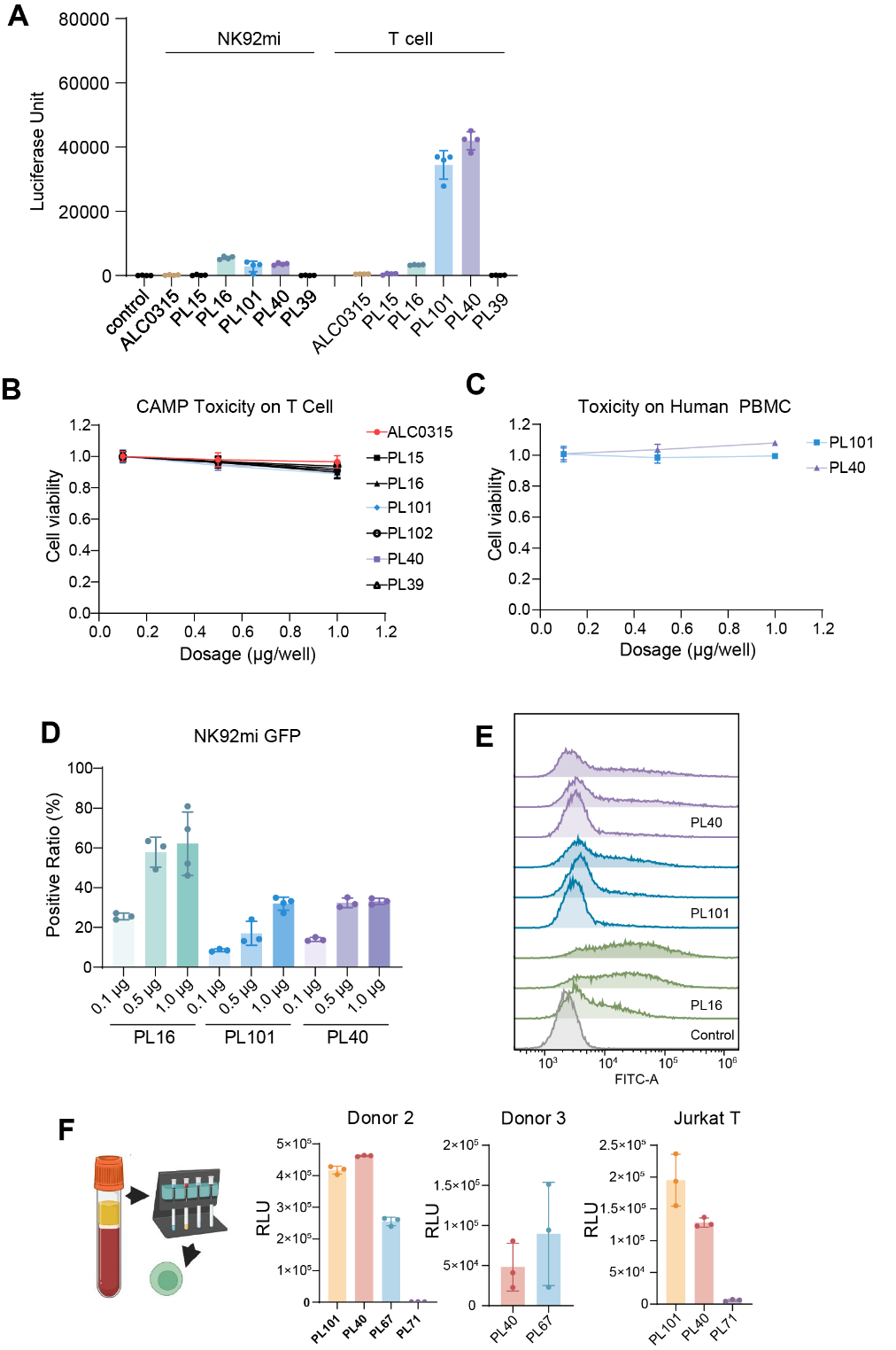

**Fig. S1. Delivery efficiency and cell viability of CAMP LNPs in multiple cell lines**. **(A)** The transfection efficiency of CAMP LNPs to various types of cells. The 96-well plates were paved with NK92mi, BMDM, A549 and human primary T cell respectively. Then, 60ng/well mLuc was transfected and measured at 24 h (n = 4). **(B and C)** The cell viability after treating with different concentrations of CAMP LNPs after 24 h. CAMP lipids were applied to transfect human primary T cell or PBMCs, at a dose of 0.1, 0.5, 1.0 μg/10^5^ cells respectively (n = 5). **(D and E)** The percentage of NK92mi cells transfected with CAMP LNPs encapsulating mGFP. 0.1 μg, 0.5 μg and 1 μg GFP mRNA were transfected to 3×10^4^ NK92mi cells, detected at 40 h using flow cytometry. **(F)** The transfection efficiency of CAMP LNPs to T cells from different human donors. 60 ng/well mLuc was transfected and measured at 24h (n = 4). Data were presented as Mean ± SD.

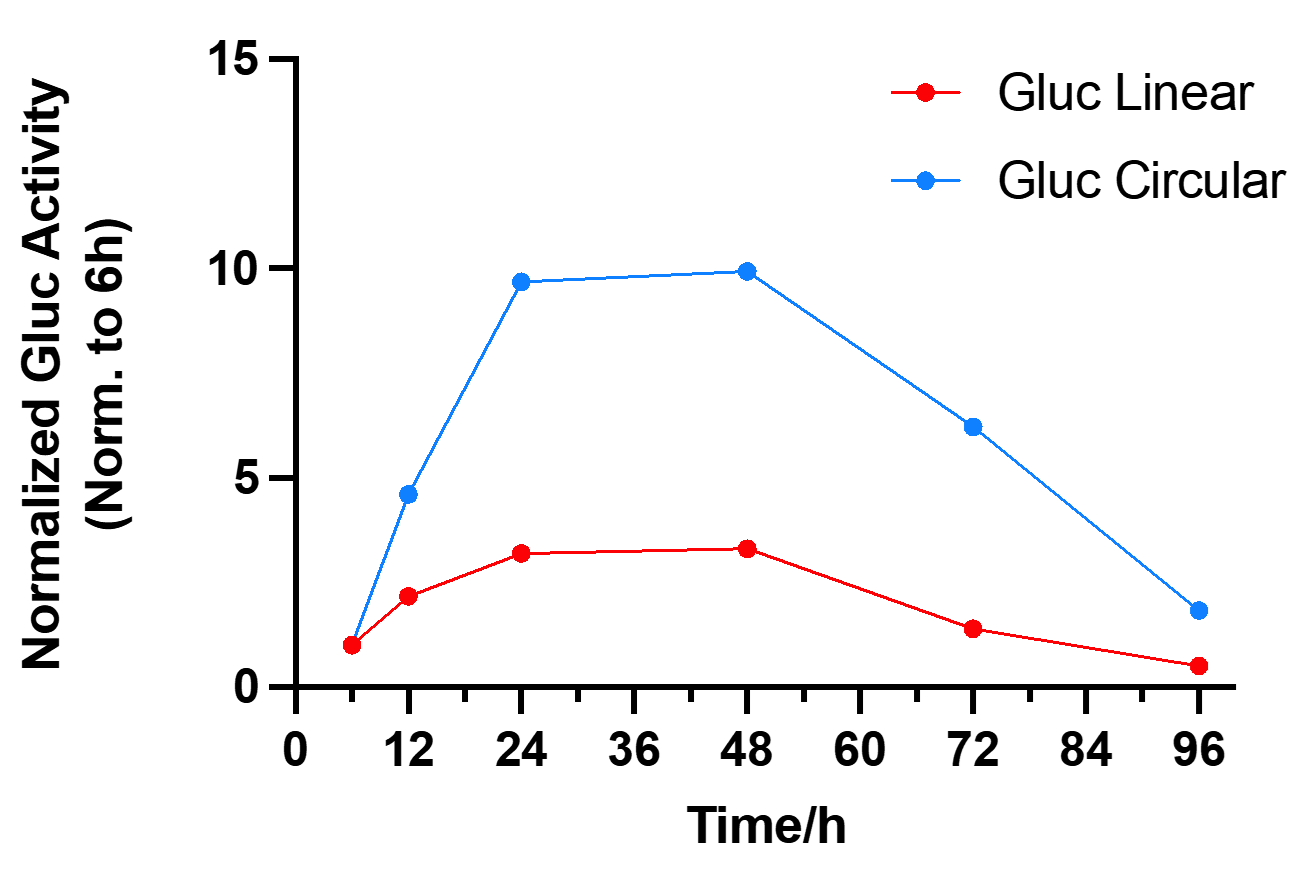

**Fig. S2. Circular RNA (Circ RNA) enhances the expression and half-life of Gluc in HEK293T cell.** HEK293T cells were plated in 96-well plates at a density of 3000 cells per well. Then, 60 ng/well circ or lin Gluc RNA was added into the well. At each detection time point, the supernatant was replaced with fresh media and Gluc was detected with coelenterazine h (n = 3), “n” indicated biologically independent samples.

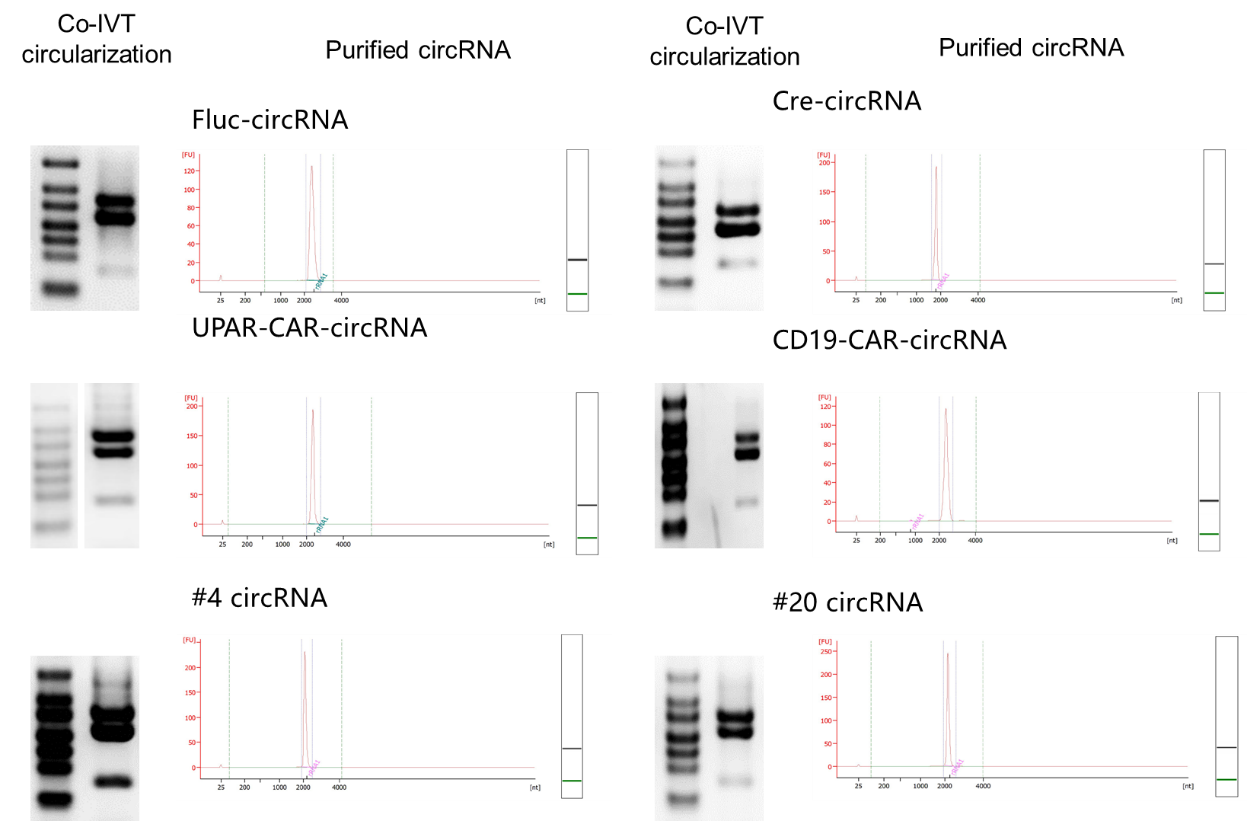

**Fig. S3. The preparation and purification of circular RNA.** The figure shows the circularization and purification with FPLC of circRNAs used in this study.

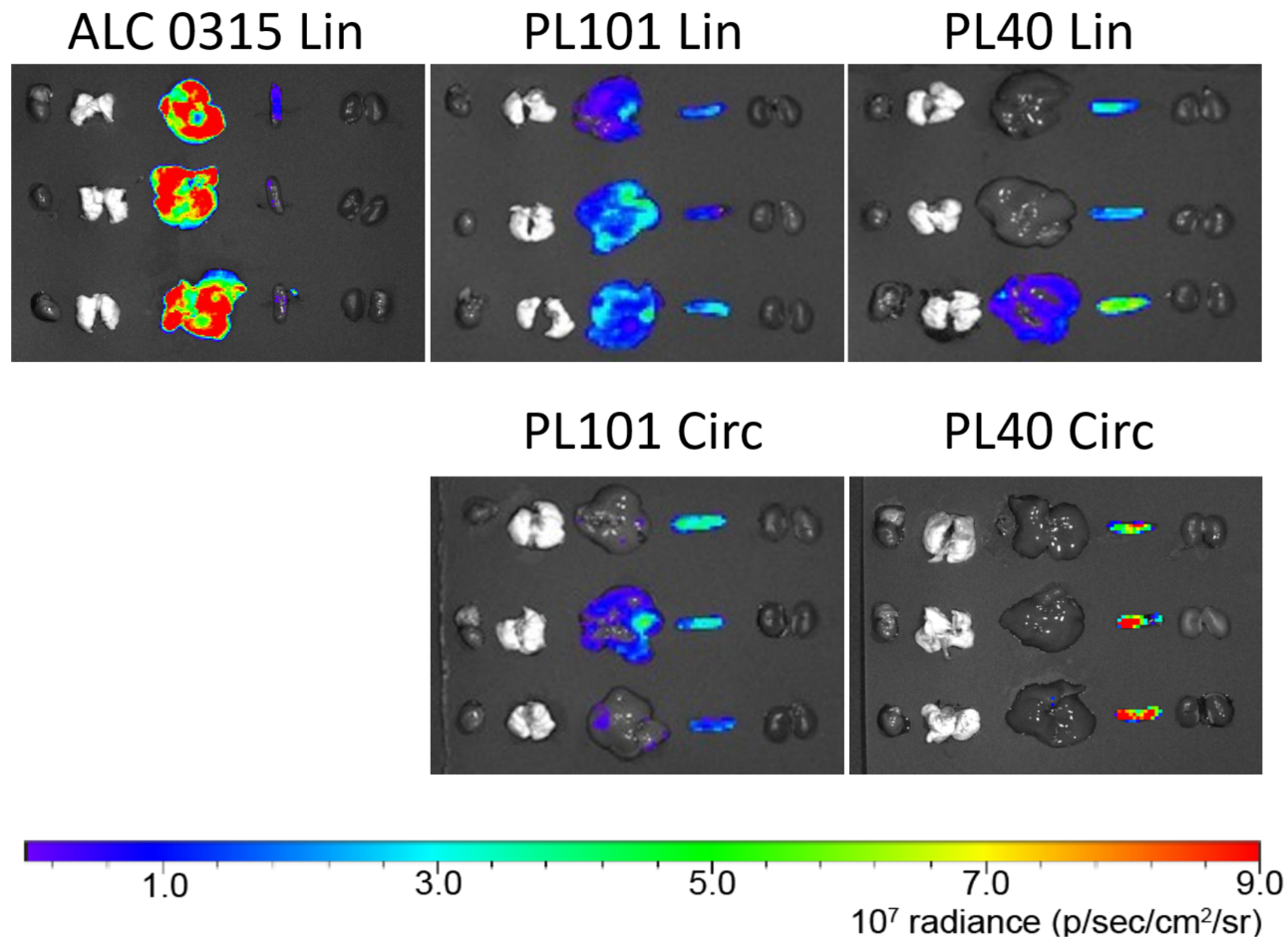
**Fig. S4. In vivo mRNA delivery efficiency with different LNPs.** IVIS images of i.v. injections of linear (Lin) or circular (Circ) mLuc mRNA LNPs. Six hours after injection, organs were collected for imaging (n = 3 biologically independent mice).

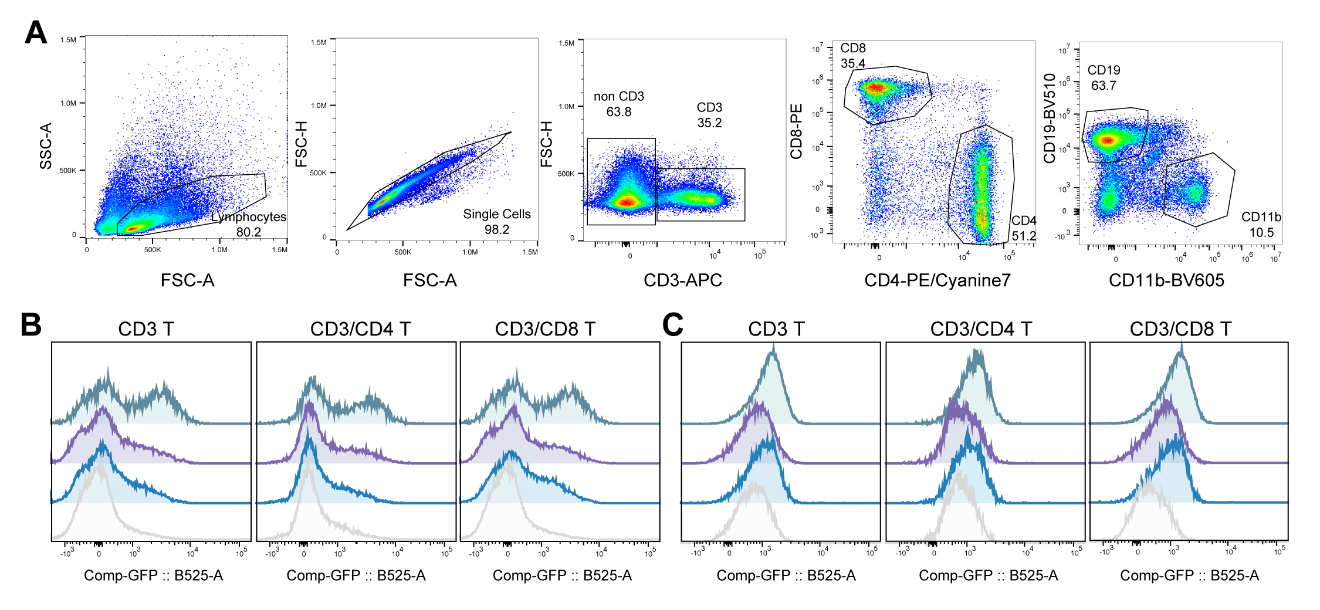

**Fig. S5. Gating strategy and T cell transfection with mCre PL40 LNP in lymph node and peripheral blood of the LoxP-GFP mice. (A)** The gating strategy of flow cytometry. **(B)** Representative histograms of T cells’ expression of GFP within lymph nodes (n = 3). **(C)** Representative histograms of T cells’ expression of GFP within peripheral blood (n = 3). “n” indicated biologically independent samples.

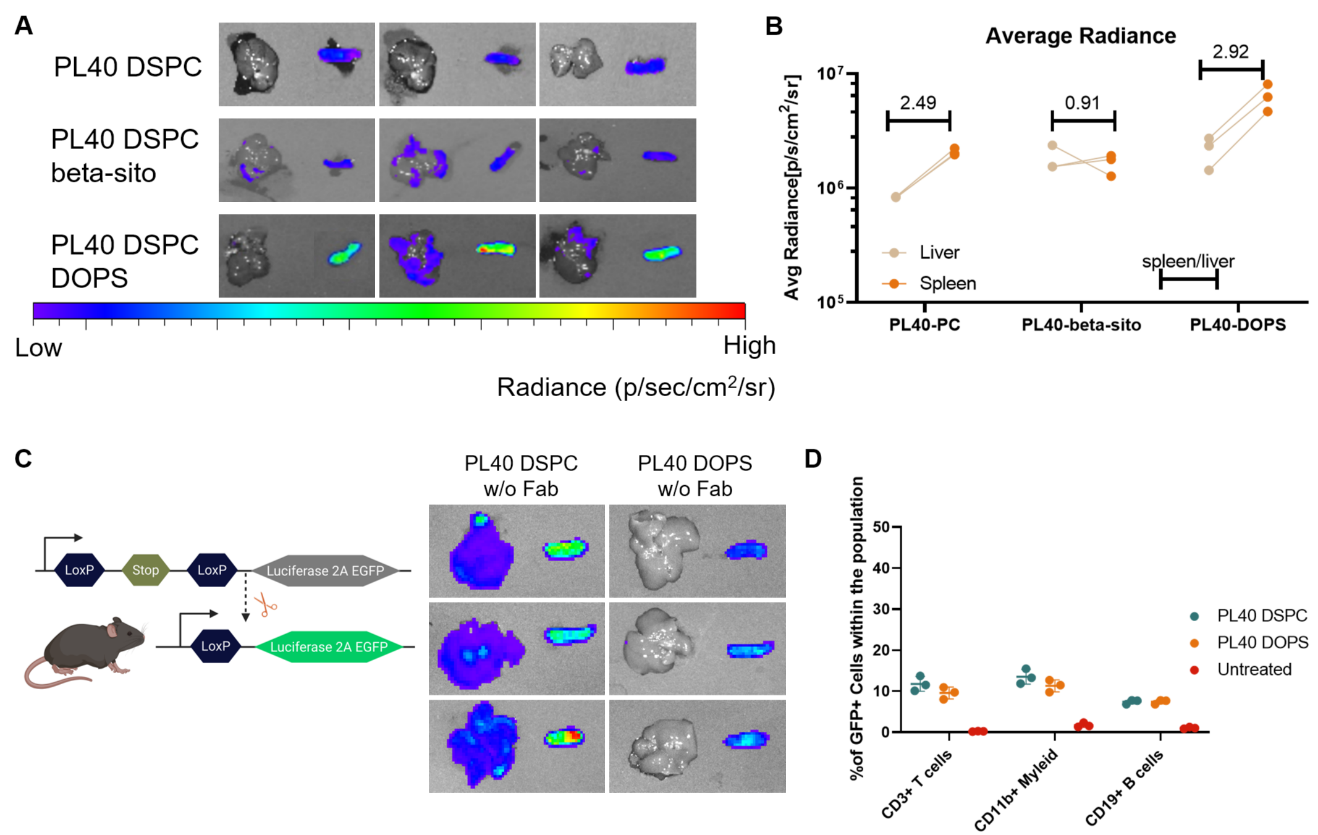

**Fig. S6. Delivery of mRNA with alternative PL40 formulations to organs and cells. (A)** IVIS images of i.v. injections of linear mLuc mRNA LNPs. Six hours after injection, organs were collected for imaging (n = 3 biologically independent mice). The LNPs were formulated with DSPC or DOPS as the helper lipids. **(B)** Quantification of expression levels of mRNA LNPs 6 hours after injection. Spleen/liver ratio was presented. **(C)** Illustration of the Cre mRNA studies in loxP-Luciferase-2A-GFP mice, and the Luciferase expression was imaged. **(D)** GFP expression in different immune cells within spleens (n = 3 biologically independent mice, data were presented as Mean ± SD).

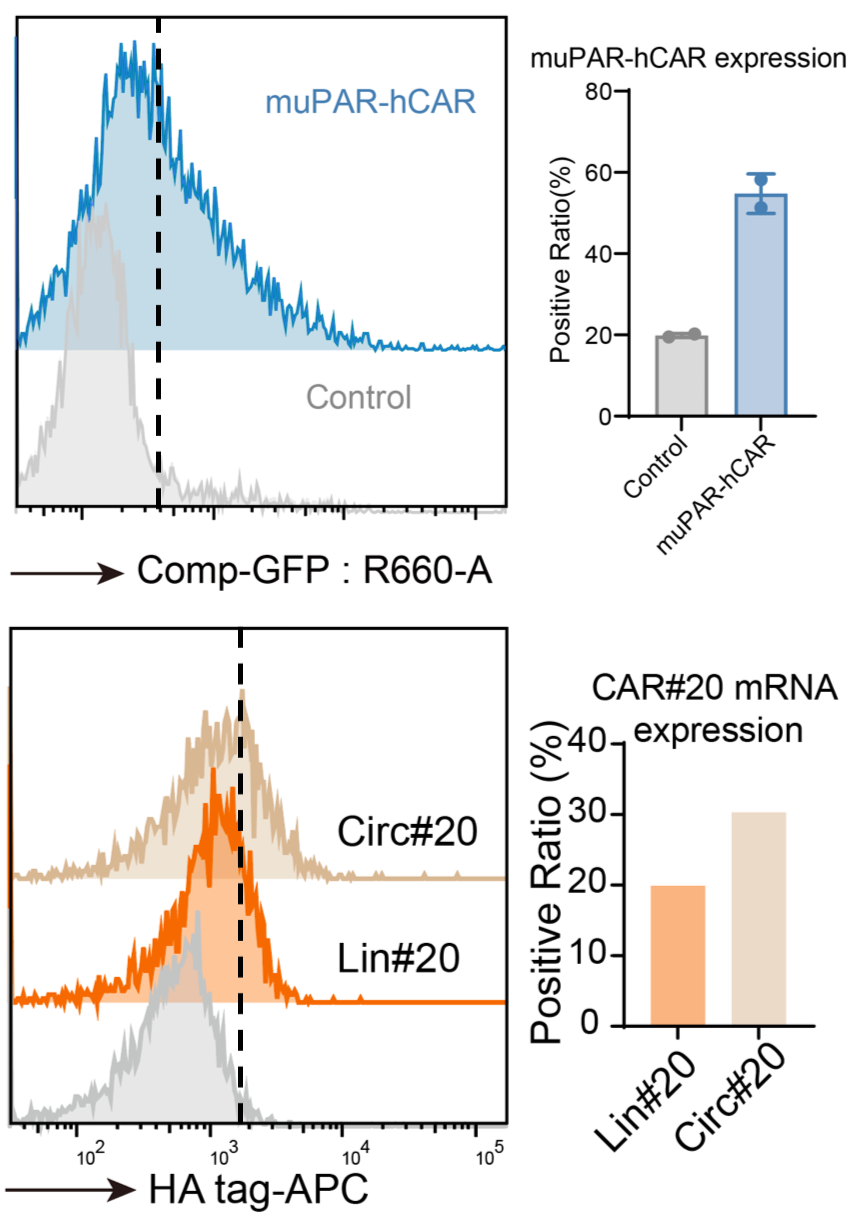

**Fig. S7. The transfection efficiency of PL40 LNP delivered muPAR-hCAR mRNA and Lin&Circ CAR#20 mRNA in primary in human T cell.** The figure shows the human primary T cell transfection results with muPAR-hCAR mRNA or circ and lin CAR#20 mRNA.

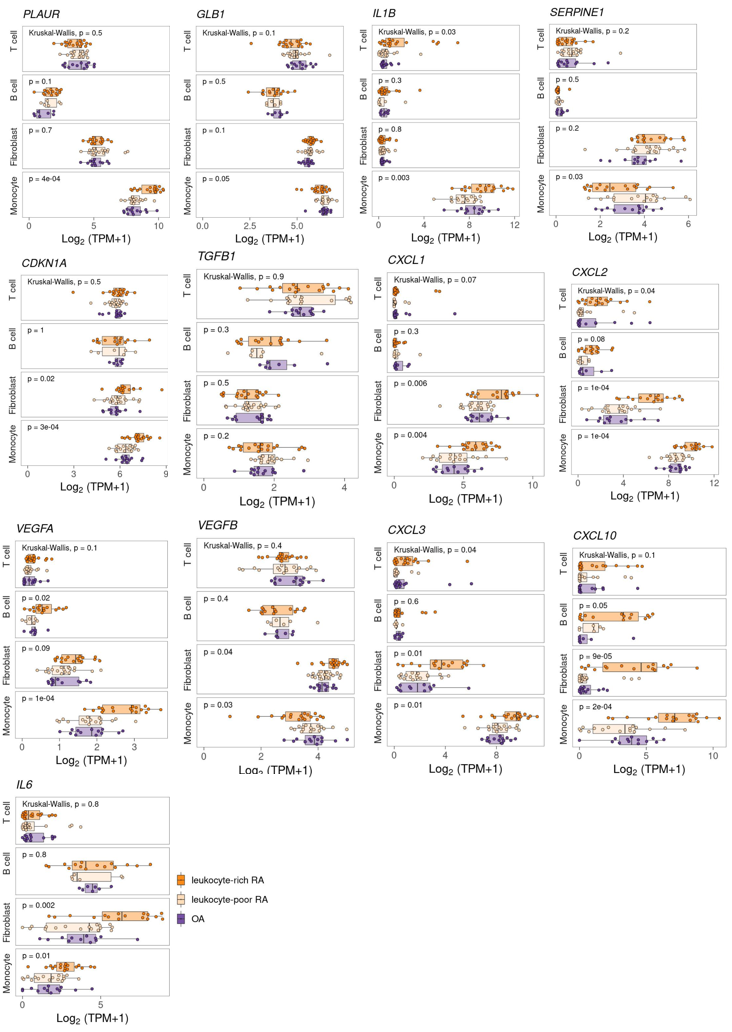

**Fig. S8. Bulk RNA-seq data of SASP and senescence marker.** The data was obtained from Immunogenomics.io, the dataset is named as AMP Phase I: Systemic Lupus Erythematosus, developed by Soumya Raychaudhuri group (*1*).

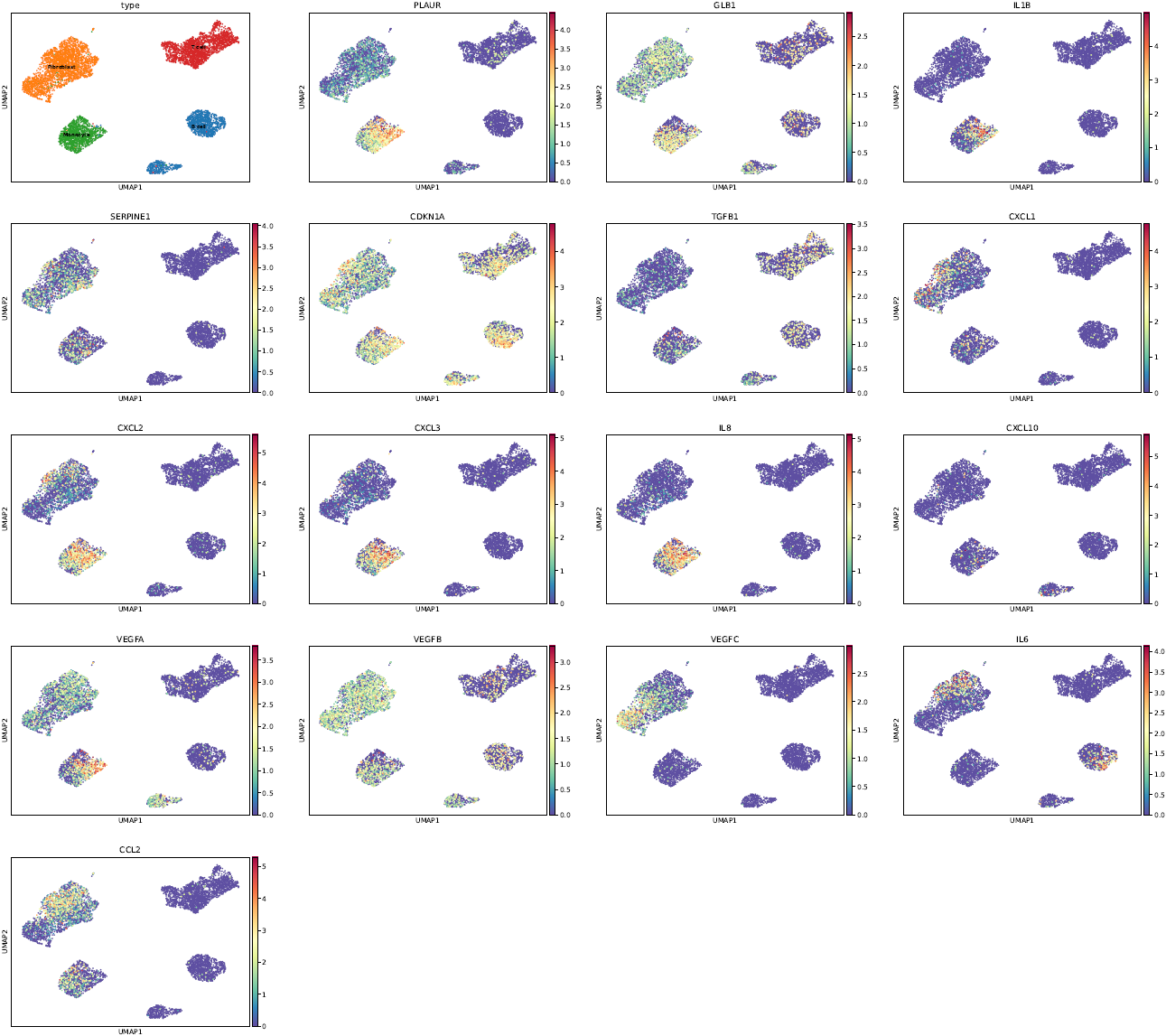

**Fig. S9. Single cell analysis of human RA data for illustration of senescence associated markers.** The scRNA-seq data were obtained from Immunogenomics.io, the dataset is named as AMP Phase I: Systemic Lupus Erythematosus, developed by Soumya Raychaudhuri group (*1*), the data was recalculated and processed.

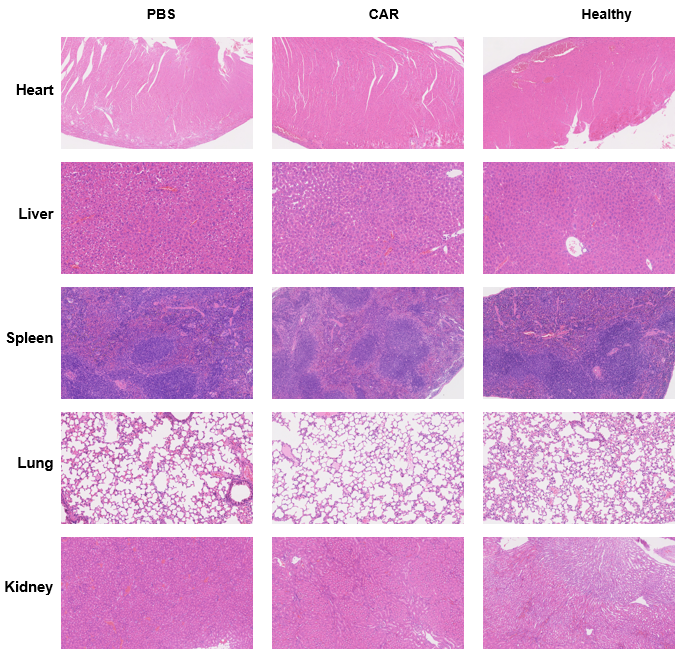

**Fig. S10. Organ toxicity of CAMP LNPs.** The main organs including heart, liver, spleen, lung and kidney from RA mice, CAR treated RA mice and healthy DBA/1 mice were collected for H&E staining. No obvious organ toxicities were observed from CAR LNP treatments. One representative image from each group is presented. The experiments were performed on 5 independent mice from each group.

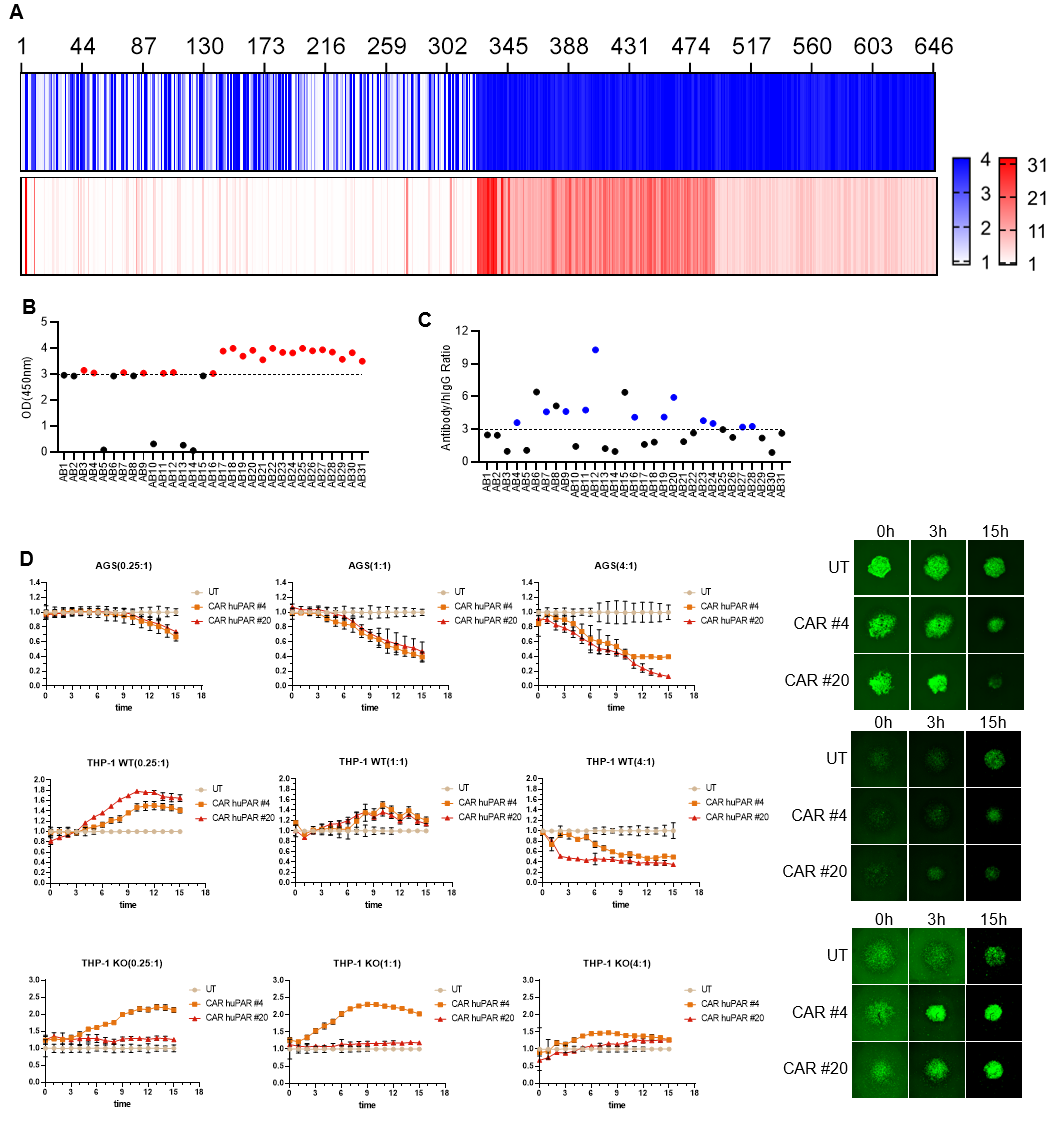

**Fig. S11. The screening process of CAR against human uPAR.** **(A)** ELISA and FACS analysis of the binding ability of the culture supernatants of 647 clones of monocolonal antibodies (mAbs) to uPAR. Thirty-one clones were obtained through comprehensive analysis. **(B and C)** mAbs with OD450nm and antibody/hIgG ratio greater than 3 were further selected from 31 clones, and 12 top-performing mAbs were obtained. **(D)** The cytotoxicity of lenti-virus transfected CAR #4 and CAR #20 were detected on two target cells and one negative cells. Human gastric adenocarcinoma AGS cell line, wild type human monocyte THP-1 cell line (THP-1 WT) and uPAR-knocked out THP-1 (THP-1 KO) were coculture with CAR #4, CAR #20 and untreated T cell, at E: T = 4: 1, 1: 1, 0.25: 1 (n = 3). CAR cargos were transfected to T cells with lenti-virus vectors. The whole cytotoxcity process was monitored by IncuCyte SX5.”n” indicates biologically independent samples.

**Supplementary Tables**

**Table S1. List of antibodies used in this study.**

| **Name** | **Reactivity** | **Application** | **Vender** | **catalog** | **Dilution** |
| --- | --- | --- | --- | --- | --- |
| PerCP/Cyanine5.5 anti-CD45 | mouse | flow cytometry(FC) | Biolegend |  | 1: 400 |
| APC/Cyanine7 anti-CD44 | mouse | FC | Biolegend |  | 1: 300 |
| PE/Cyanine7 anti-CD62L | mouse | FC | Biolegend |  | 1: 300 |
| APC anti-CD3 | mouse | FC | Biolegend |  | 1: 400 |
| FITC anti-CD4 | mouse | FC | Biolegend |  | 1: 400 |
| PE anti-CD8 | mouse | FC | Biolegend |  | 1: 400 |
| BV421 anti-PD1 | mouse | FC | Biolegend |  | 1: 300 |
| APC/Cyanine7 anti-CD45 | mouse | FC | Biolegend |  | 1: 400 |
| BV605 anti-CD11b | mouse | FC | Biolegend |  | 1: 400 |
| APC anti-CD11c | mouse | FC | Biolegend |  | 1: 300 |
| BV510 anti-F4/80 | mouse | FC | Biolegend |  | 1: 400 |
| BV421 anti-CD86 | mouse | FC | Biolegend |  | 1: 300 |
| PE/Cyanine5 anti-CD206 | mouse | FC | Biolegend |  | 1: 300 |
| FITC anti-CD3 | mouse | FC | Biolegend |  | 1: 400 |
| PerCP/Cyanine5.5 anti-CD4 | mouse | FC | Biolegend |  | 1: 400 |
| APC anti-CD25 | mouse | FC | Biolegend |  | 1: 400 |
| PE/Cyanine7 anti-CD69 | mouse | FC | Biolegend |  | 1: 300 |
| BV421 anti-Foxp3 | mouse | FC | Biolegend |  | 1: 300 |
| BV510 anti-IFNγ | mouse | FC | Biolegend |  | 1: 300 |
| BV510 anti-CD19 | mouse | FC | Biolegend |  | 1: 300 |
| APC-HA.11 epitope tag |  | FC | Biolegend |  | 1: 200 |
| anti-αSMA | mouse | mIHC | CST | 14968S | 1: 1000 |
| anti-F4/80 | mouse | mIHC | CST | D2S9R | 1: 500 |
| anti-HA tag |  | mIHC | CST | C29F4 | 1: 1000 |
| anti-CD206 | mouse | mIHC | Abcam | AB300621 | 1: 2000 |
| anti-CD3ε | mouse | mIHC | CST | E4T1B | 1: 400 |
| anti-uPAR | mouse | mIHC | Abcam | AB307895 | 1: 100 |
